# Advancing chloroplast synthetic biology through high-throughput plastome engineering of *Chlamydomonas reinhardtii*

**DOI:** 10.1101/2024.05.08.593163

**Authors:** René Inckemann, Tanguy Chotel, Cedric K. Brinkmann, Michael Burgis, Laura Andreas, Jessica Baumann, Priyati Sharma, Melanie Klose, James Barrett, Fabian Ries, Nicole Paczia, Timo Glatter, Felix Willmund, Luke C. M. Mackinder, Tobias J. Erb

**Author notes:** These authors contributed equally to this work (René Inckemann, Tanguy Chotel). Corresponding authors Tobias J. Erb, Max-Planck Institute for Terrestrial Microbiology, Marburg 35043, Germany; Center for Synthetic Microbiology, Philipps-Universität Marburg, Marburg 35032, Germany.

## Abstract

Chloroplast synthetic biology holds promise for developing improved crops through improving the function of plastids. However, chloroplast engineering efforts face limitations due to the scarcity of genetic tools and the low throughput of plant-based systems. To address these challenges, we here established *Chlamydomonas reinhardtii* as a prototyping chassis for chloroplast synthetic biology. We developed an automation workflow that enables the generation, handling, and analysis of thousands of transplastomic strains in parallel, expanded the repertoire of selection markers for chloroplast transformation, established new reporter genes, and characterized over 140 regulatory parts, including native and synthetic promoters, UTRs, and intercistronic expression elements. We integrated the system within the Phytobrick cloning standard and demonstrate several applications, including a library-based approach to develop synthetic promoter designs in plastids. Finally, we provide a proof-of-concept for prototyping novel traits in plastids by introducing a chloroplast-based synthetic photorespiration pathway and demonstrating a twofold increase in biomass production. Overall, our study advances chloroplast engineering, and provides a promising platform to rapidly prototype chloroplast manipulations before their transfer into higher plants and crops.

---

Synthetic biology offers promising prospects for developing photosynthetic organisms and crop varieties with improved traits.^1^ These include enhanced resilience to extreme weather conditions, such as heat waves and drought^2^, superior nutrient content, like omega fatty acids^3^ and vitamins^4^, and improved yield, for instance through more efficient photorespiratory bypasses^5–10^ or entirely new-to-nature carbon fixation cycles.^11–15^ However, realizing the full potential of synthetic biology in photosynthetic eukaryotes will require significant advancements in genetic engineering capabilities that go beyond traditional breeding and gene-editing techniques.^16^

In particular chloroplasts have become an increasing target for advanced engineering efforts.^7,17–19^ These organelles harbor the photosynthetic apparatus and house several metabolic processes of high interest for the engineering of novel traits. For example chloroplast genome (plastome)-based engineering of plastid isoprenoid metabolism has recently proven a highly effective strategy for the production of high-value compounds, such as astaxanthin or artemisinin.^20,21^ Moreover, several subunits of the light-harvesting and carbon fixation machineries are encoded by the plastome, which makes the direct manipulation of the chloroplast genome central for any efforts that aim at improving photosynthetic yield. One current example are efforts of integrating carboxysome and pyrenoid bases CO2-concentrating mechanism into plant chloroplasts to boost photosynthetic efficiency^22–30^

These examples emphasize the growing need to develop advanced genetic tools for the direct manipulation of the plastome. However, chloroplast genome engineering offers additional advantages compared to nuclear genome-based engineering strategies of chloroplast functions. Expression of transgenes from the chloroplast genome benefits from precise genomic integration, the absence of (nuclear) silencing mechanisms, significant higher protein levels that can be achieved, and improved genetic containment, as transgenes are not transmitted by pollen due to the strict maternal inheritance of chloroplasts.^31^

Yet, despite all these advantages, genetic engineering of the plastid is still constrained to only a handful of available genetic elements and tools.^32,33^ Currently, only one or two selection markers are used on a routine base.^34^ Moreover, only a small number of natural gene expression elements (5’UTR, 3’UTR, promoters) are available, which limits the number of genes that can be stacked, and provides little control over their expression strength.^32^ This limited tool set is even more constrained, as gene expression elements cannot be simply reused because of the high homologous recombination frequency of the plastome, which is observed for sequences as short as 50 base pairs.^35^

Thus, to bring the applications of synthetic biology in plastids to the level of versatility and complexity that is available for other organisms, methods for the systematic assembly and large-scale characterization of gene expression elements in chloroplasts are urgently required.^19,36,37^ However, the long generation times of photosynthetic eukaryotes, and the absence of suitable model systems that are compatible with high-throughput handling routines have impeded the development of genetic tools for the chloroplast, thus far.^38^ Consequently, the systematic development of plastid transgene expression strategies and the prototyping of complex genetic designs in chloroplasts have remained beyond reach.

In this study, we aimed at overcoming above shortcomings, by establishing *Chlamydomonas reinhardtii* as chassis for advancing chloroplast synthetic biology. This fast-growing, unicellular green algae contains a single chloroplast and can be cultivated in high-throughput formats^39–42^, which makes it a highly suitable model for plastome-engineering. To generate, handle and analyze thousands of transplastomic *C. reinhardtii* strains in parallel, we developed an automation workflow, expanded the number of selection markers for chloroplast transformation, and established several new reporter genes for fluorescence and luminescence-based readouts and cell sorting.

Building on these resources, we systematically characterized a collection of more than 140 regulatory parts, including 35 different 5’UTRs, 36 3’UTRs, 59 promoters, and 16 intercistronic expression elements (IEEs) for advanced gene stacking. We embedded these genetic parts within a Phytobrick/Modular cloning (MoClo) framework^19^ for the standardized and automated assembly of genetic constructs that are compatible with existing *C. reinhardtii*, and plant MoClo resources. Using this MoClo routine allowed us to rapidly assemble and characterize gene element combinations at multiple insertion loci to establish multi-transgene constructs that range across more than three orders of magnitude in expression strength. Finally, we showcase the utility of our tools by developing more than 30 synthetic promoters through a pooled library-based approach in chloroplasts and provide another real-world application case by introducing a recently reported synthetic photorespiration pathway into the chloroplast^7^.

Overall, our work provides a high-throughput workflow for chloroplast synthetic biology and a foundational set of genetic elements for the convenient engineering of the chloroplast that is freely available to the community. Because of a high transferability of results between the chloroplasts of *C. reinhardtii* and the chloroplasts of other organisms^43–45^, our platform opens the possibility for rapid prototyping of new traits before transfer into higher plants and crops.

## Results

### Automation workflow for high-throughput characterization of transplastomic *C. reinhardtii* strains

To facilitate the systematic characterization of genetic parts across thousands of transplastomic *C. reinhardtii* strains, we developed an automated chloroplast engineering workflow. This workflow is based on solid-media cultivation and leverages a contactless liquid-handling robot to enhance our capacity for handling an increased number of strains (Fig. 1). In brief, the workflow relies on the automated picking of transformants into a standardized 384 format and subsequent restreaks to achieve homoplasy, using a Rotor screening robot. These colonies are then organized into a 96-array format for high-throughput biomass growth, liquid-medium transfer and/or additional screens such as reporter gene analysis. The later involves using the Rotor screening robot for the transfer of biomass from 96-array agar plates into multi-well plates filled with water. Following resuspension, the optical density is measured at 750 nm (OD750), and a contact-free liquid handler is used for cell number normalization, medium transfer, and the supplementation of additional compounds (e.g., the NanoLuc substrate). Transitioning to a (partially) solid-media based workflow proved to be an advancement over previous techniques that solely relied on liquid-medium cultures, which inherently limited the number of strains that can be handled in parallel. Overall, this automation workflow was essential for generating and managing the 3,156 individual transplastomic strains that were used in this study, as presented in the following.

**Figure 1:**
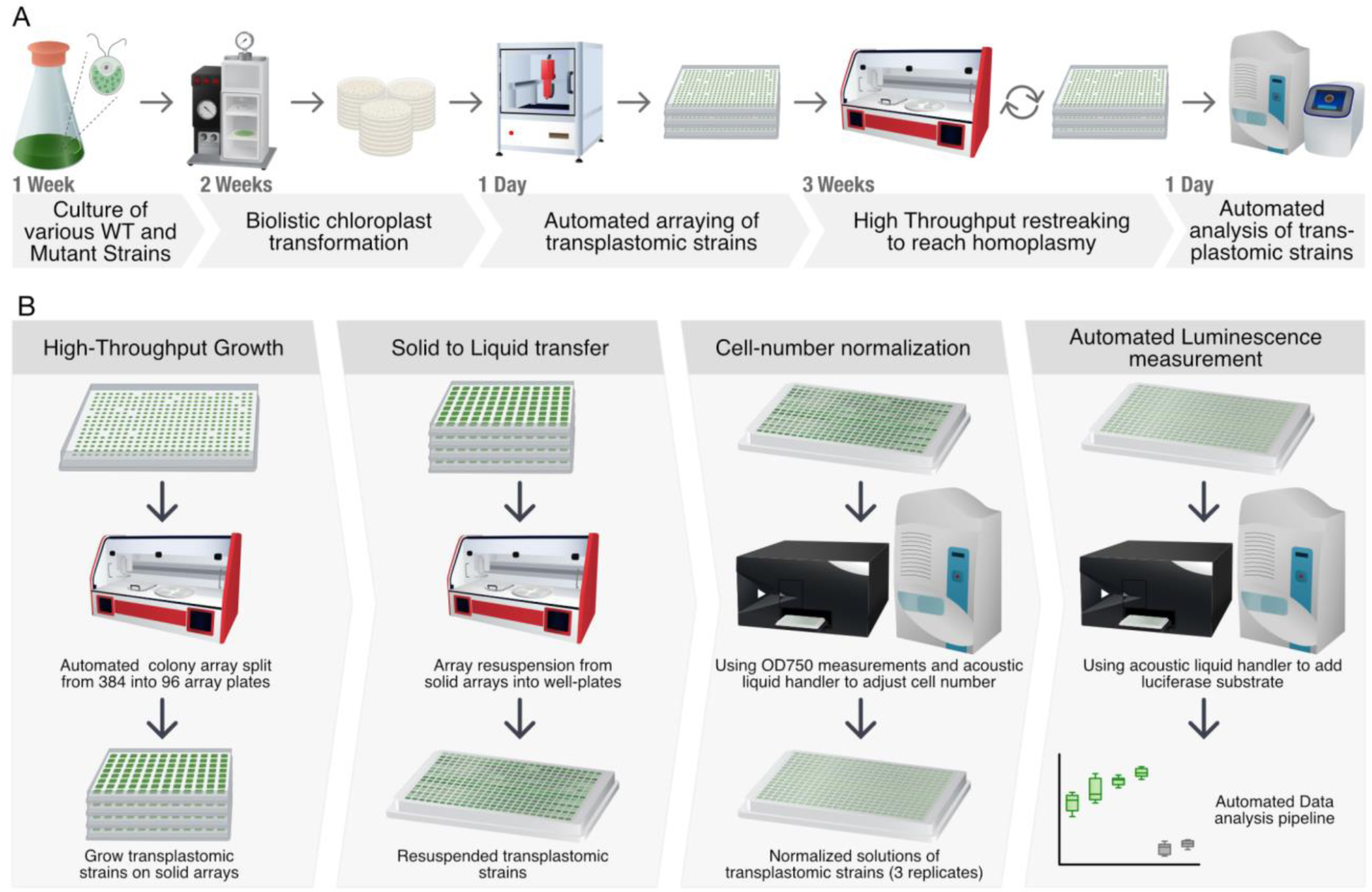
Workflow for generation and high-throughput characterization of transplastomic *C. reinhardtii* strains. **a,** Automated transplastomic strain generation workflow. Chloroplast transformation is performed via biolistic transformation via particle gun, subsequent automated arraying of transformants via PixL picking robot. Transplastomic strains are restreaked three times to reach homoplasmy via Rotor screening robot. Automated genotyping of transplastomic strains is performed via acoustic liquid handler and qPCR analysis to confirm homoplasmy. **b** High throughput characterization of transplastomic strains. Colonies are transferred to 96-well formats for biomass cultivation, preparing for reporter gene measurement. Rotor robot is then used to transfer biomass from agar (96 strains/plate) to multiwell plates for resuspension. Optical density at 750 nm (OD_750_) is measured, and the Echo liquid handler is used for cell normalization and NanoLuc luciferase substrate addition.

### Establishing a foundational set of genetic parts for chloroplast engineering

We next established a foundational set of >300 genetic parts for plastome manipulation of *C. reinhardtii*, which we embedded in a standardized MoClo framework^46,47^ (Fig. 2a, Extended Data Fig. 1). This set of parts encompasses regulatory elements, such as 5’UTRs, 3’UTRs, Intercistronic expression elements (IEEs), derived from the chloroplast genome of *C. reinhardtii*, tobacco and even includes some synthetic regulatory elements. Additionally, it also contains parts for integration into various sites within the chloroplast genome (Extended Data Fig. 1). Our repository also encompasses several selection markers and reporter genes, which we identified and validated for *C. reinhardtii*. Beyond the well-established spectinomycin selection marker, we introduced tobramycin as a selection system for chloroplast transformation in *C. reinhardtii* (Fig. 2b, Extended Data Fig 2.c). This selection marker exhibited transformations efficiencies that are comparable to spectinomycin-based plastome selection, with 100-1,000 colonies per plate and ∼10µg DNA biolistic transformation. We also integrated and validated kanamycin^48^ and phosphite^49,50^ selection markers in our parts library, as well as an arginine-based selection system^51^ that complements arginine auxotrophy in a *C. reinhardtii* Arg9 deletion background (Fig. 2b, Extended Data Fig. 2d).

**Figure 2:**
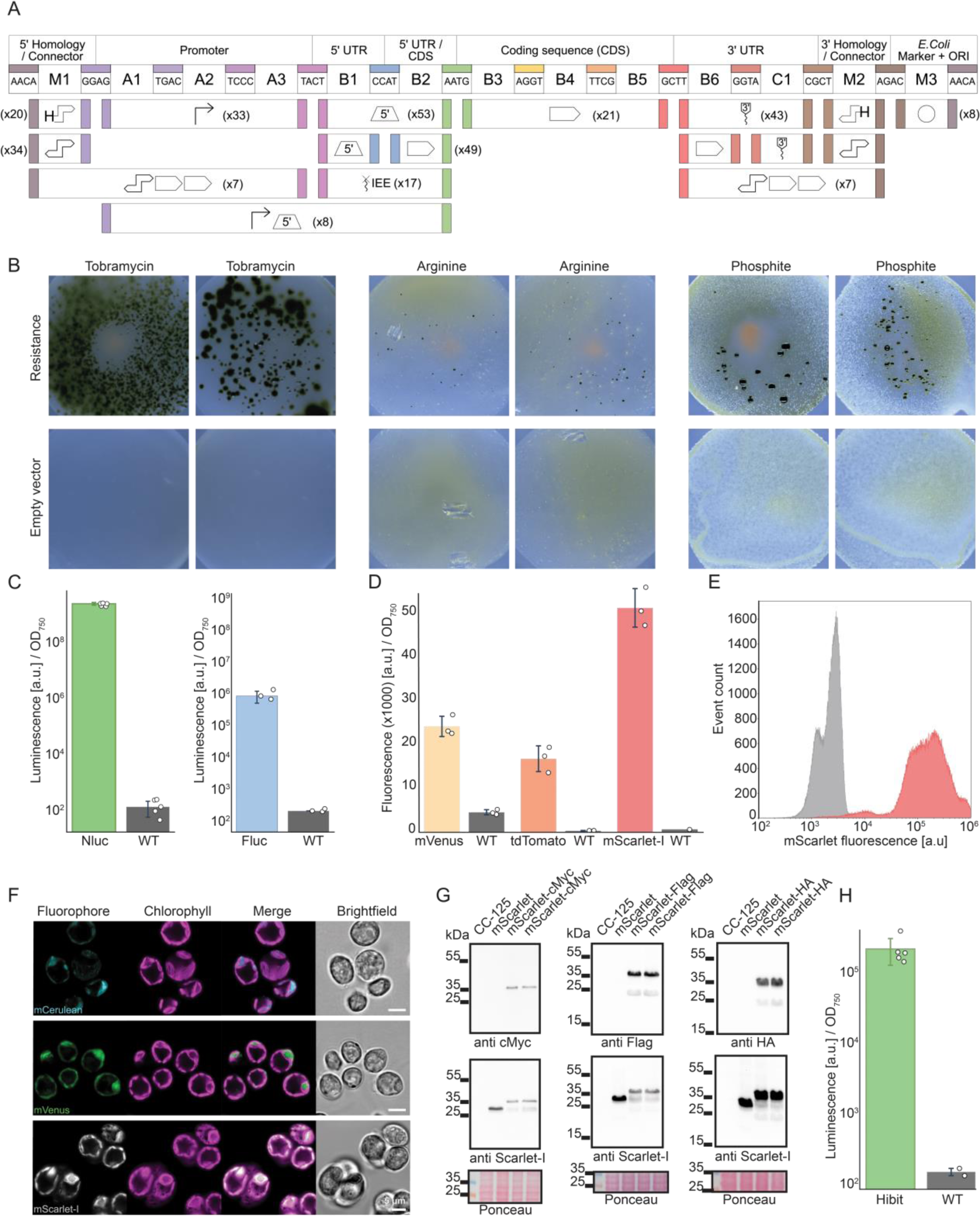
Establishing novel basic tools for chloroplast engineering. **a,** Architecture of the novel chloroplast modular cloning (MoClo) system and overview of the created genetic parts. The architecture follows the Phytobrick standard^47^ and conserves the cloning overhangs of this system, as well as the nomenclature for the part types (A1-C1). The position M1-3 allow for higher modularity as reported by Stukenberg *et al.* Number of parts are indicated for each position **b,** Selection markers for chloroplast transformation. Three different selection marker systems are shown with two replicates and two negative controls for each one. Successful selection after chloroplast transformation can be observed for all three of them, notably with a high efficiency for tobramycin-based selection (100µg/mL). **c,** NanoLuc as a luminescence reporter for the chloroplast. Nanoluc signal of a transplastomic strain containing a NanoLuc expression cassette (green) is plotted as arbitrary units [a.u.] normalized to OD and compared to WT strain (gray). **d,** Novel fluorescent reporters for the chloroplast. Fluorescent signals of transplastomic strains containing different fluorescent reporter expression cassettes, measured via a plate reader, are plotted as arbitrary units [a.u.] normalized to OD_750_ and compared to WT strain (gray). **e,** Flow cytometry data of the transplastomic strain, containing a mScarlet-I expression cassette (red) compared to the WT strain (gray). **f,** Fluorescence microscopy images of the transplastomic strains containing different fluorophores (mCerluean, mVenus and mScarlet-I). **g,** Western blot analysis of transplastomic strains producing mScarlet FLAG-tagged, HA-tagged, cMyc-tagged in the C-term position. Each strain is also tested for the presence of mScarlet using mScarlet-I specific probes. Ponceau gels are shown for each gel. **h,** Nanoluc assay for the Hibit tag in C-term position. Data is plotted as arbitrary units [a.u.] normalized to OD_750_ and compared to WT strain (gray).

We also tested different luciferase-based and fluorescent reporters for the characterization and quantification of regulatory elements. As primary reporter, we identified NanoLuc luciferase (Extended Data Fig. 2a). This luminescence reporter did not interfere with chloroplast auto-fluorescence, and showed a high signal-to-noise ratio that was superior to that of established fluorescent reporters (Fig. 2c and 2d). Notably, it produced a luminescence signal that was nearly seven orders of magnitude higher than that of the wild-type background (Fig. 2c). As alternative to NanoLuc, we integrated the previously reported firefly luciferase^52^ into our parts library. Furthermore, we tested four different fluorescent reporters (mVenus, mScarlet-I, td tomato and mCerulean), of which mScarlet-I performed best. This reporter showed little interference with chloroplast auto-fluorescence (Fig. 2d) and could be successfully applied in flow cytometry and microscopy (Fig. 2f). While the fluorescence of mCerulean measured in the plate reader was only 1.5 times higher (Extended Data Fig. 2b) than the wild-type signal on average, it was still sufficient to facilitate effective microscopy analysis (Fig. 2f).

Finally, we included additional parts for gene product detection into our library, which included modular tags (FLAG-, HA, cMyc-tags) for western blots and a HiBiT-luciferase complementation assay to tag and detect proteins of interest (Fig. 2g, h, Extended Data Fig2.e). This set of selection markers, reporter genes and regulatory parts provided the basis for all high-throughput characterization efforts and engineering applications of this study.

### High-throughput characterization of regulatory elements

Having established the initial library of parts, we proceeded to systematically quantify the different regulatory parts using the NanoLuc based read-out. To that end, we tested various 5’ untranslated regions (5’UTRs), as well as 3’UTRs, which both are known to significantly influence and regulate gene expression in the chloroplast.^53–56^ Despite their cyanobacterial origins, chloroplasts diverge from their prokaryotic ancestors in gene expression regulation.^57^ While prokaryotic gene expression is predominantly controlled at the transcriptional level, chloroplast gene expression is mainly regulated post-transcriptionally via RNA elements in the 5’ and 3’ UTRs.^58^ Especially 5’UTRs are known to interact with chloroplast mRNA binding factors that regulate transcript maturation, stability and translation efficiency, thus determining gene expression levels.^59–63^

To test 5’UTRs, we constructed 35 different genetic constructs, each composed of the *rrn16* promoter, followed by a variable 5’UTR, the NanoLuc luciferase coding sequence, the *psbA* 3’UTR, and homology sequences for integration at the psbH locus, which is the most widely used integration site in *C. reinhardtii*. As 5’UTR, we used natural sequences sourced from *C. reinhardtii* and tobacco, as well as synthetic sequences. For each of the 35 designs, we picked 16 biological replicates, which we tested on two different growth media (TAP and HSM) and two different light conditions (light and dark). In total, this experiment involved the handling and analysis of 2,240 transplastomic strains.

The different transplastomic strains showed a broad luminescence, which spanned over more than three orders of magnitude (Fig. 3a). Strains carrying 5’UTRs of RNA polymerase and ribosomal proteins (*rpoC2, rlp16, rpl23*) showed on average lower expression levels, while strains carrying 5’UTRs of highly abundant proteins in photosynthesis (*rbcL*, *psaA, psbC*) showed high expression levels, aligning well with recently reported Ribo-seq data for Chlamydomonas.^64^ These expression patterns proved to be consistent across the four different growth conditions, with the exception of the *petD* 5’UTR that somehow displayed divergent expression patterns across the two different media (Fig. 3b, (Extended Data Fig3.a-c).

**Figure 3:**
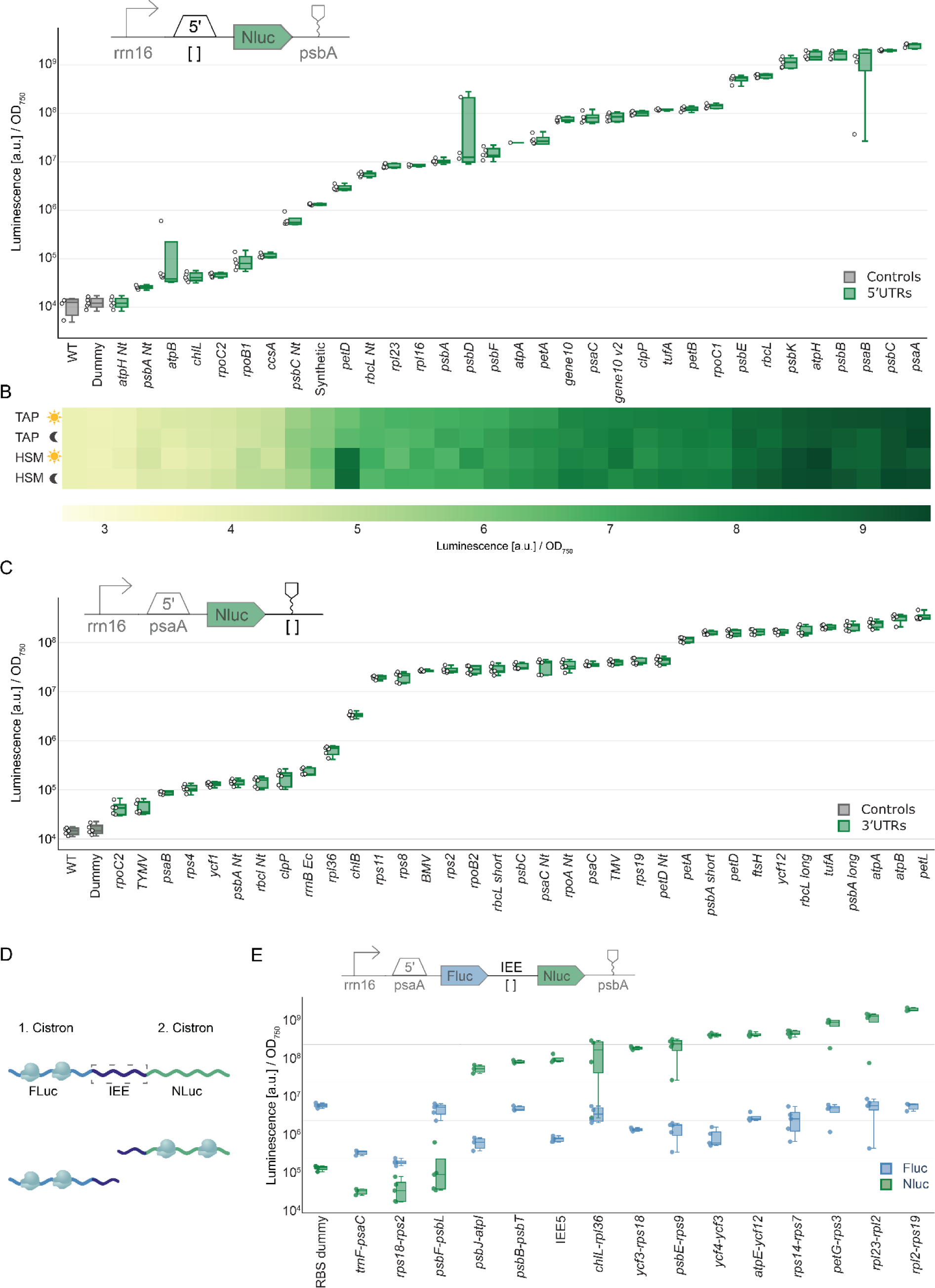
Characterization of regulatory parts for chloroplast gene expression. **a,** 5’UTR characterization 5’UTRs were characterized by measuring NanoLuc activity of transplastomic strains containing different constructs, which differ in their 5’UTR. Nanoluc signal of a transplastomic strain containing a NanoLuc expression cassette (green) is plotted as arbitrary units [a.u.] normalized to OD and compared to WT strain (gray). For all measurements, n_biological_= 5, n_technical_= 3. **b,** 5’ characterization under four different conditions. Heatmap is showing the Nanoluc signal of transplastomic strains under light, dark, TAP and HSM medium conditions. Similar expression patterns can be observed under all four conditions. **c,** 3’UTR characterization. 3’UTRs were characterized by measuring NanoLuc activity of transplastomic strains containing different constructs, which differ in their 3’UTR. Nanoluc signal of a transplastomic strain containing a NanoLuc expression cassette (green) is plotted as arbitrary units [a.u.] normalized to OD_750_ and compared to WT strain (gray). For all measurements, n_biological_= 5, n_technical_= 3 **d,** Function of Intercistronic expression elements (IEEs). Two cistrons are connected via an IEE. After processing via imported cis-factors, both cistrons are translated separately. **e,** IEE characterization. IEE characterization. IEEs were characterized by measuring NanoLuc and Fluc activity of transplastomic strains containing different constructs, which differ in their IEE. Luciferase signal of a transplastomic strain containing a NanoLuc expression cassette (green) is plotted as arbitrary units [a.u.] normalized to OD750 and compared to Fluc luminescence (blue). The dotted line differentiates IEEs with insufficient support for robust expression (left) from those facilitating strong expression (right). For all measurements, n_biological_= 5, n_technical_= 3.

Next, we focused on 3’UTRs. We constructed 36 different genetic constructs targeting the psbH locus, each carrying the *rrn16* promoter, the *psaA* 5’UTR, the NanoLuc luciferase coding sequence, and a variable 3’UTR, which we derived from *C. reinhardtii*, tobacco, or viral origin. Luminescence of the different transplastomic strains showed three levels of variability spanning across three orders of magnitude (Fig. 3c), low, medium and high, illustrating the relevance of 3’UTR on gene expression activity in chloroplasts, although to a lesser extent compared the 5’UTR.

Finally, we investigated intercistronic expression elements (IEEs) that are crucial for polycistronic gene expression in the chloroplast. IEEs promote processing of polycistronic mRNAs into stable monocistronic transcripts and protect these mRNAs against degradation^65,66^ (Fig. 3d). Utilizing small RNA profiling data from the chloroplast of Chlamydomonas^20^, we identified 15 potential candidates, which we tested as IEE between Firefly luciferase and NanoLuc luciferase as reporter genes (Fig. 3e). With exception of three sequences (*trnF-psaC; rps18-rps2; psbF-psbL*) all other sequence indicated successful processing into monocistronic mRNAs. Interestingly, the different functional IEEs showed up to tenfold variation in luciferase activity levels, highlighting the potential of these elements in modulating the expression of stacked transgenes within the chloroplast (Fig. 3e). In summary, these experiments established and quantified the behavior of more than 80 genetic elements in the chloroplast.

### Developing synthetic chloroplast expression elements through rational and library-based approaches

Most of the genetic elements tested thus far were of natural origin. Thus, we sought in the following to establish fully synthetic (i.e., non-native) genetic parts, in particular promoters. Inspired by the architecture of the *C. reinhardtii rrn16* promoter, we developed a minimal scaffold, consisting of only 46 base pairs carrying canonical −35 and −10 motifs. We preserved these −35 and −10 motifs across the majority of these promoters and modified the rest of the sequence to modulate expression strength and reduce the risk of unintended homologous recombination for multi-gene constructs.

In total, we rationally designed 21 different promoters and tested them in a standardized construct that was integrated at the psbH locus and included the different promoter sequences upstream of the psaA 5’UTR, the coding sequence of NanoLuc luciferase, and the psbA 3’UTR. The luminescence signal of the transplastomic strains varied across three orders of magnitude, suggesting that our synthetic promoters covered a range of different expression strengths (Fig. 4a).

**Figure 4:**
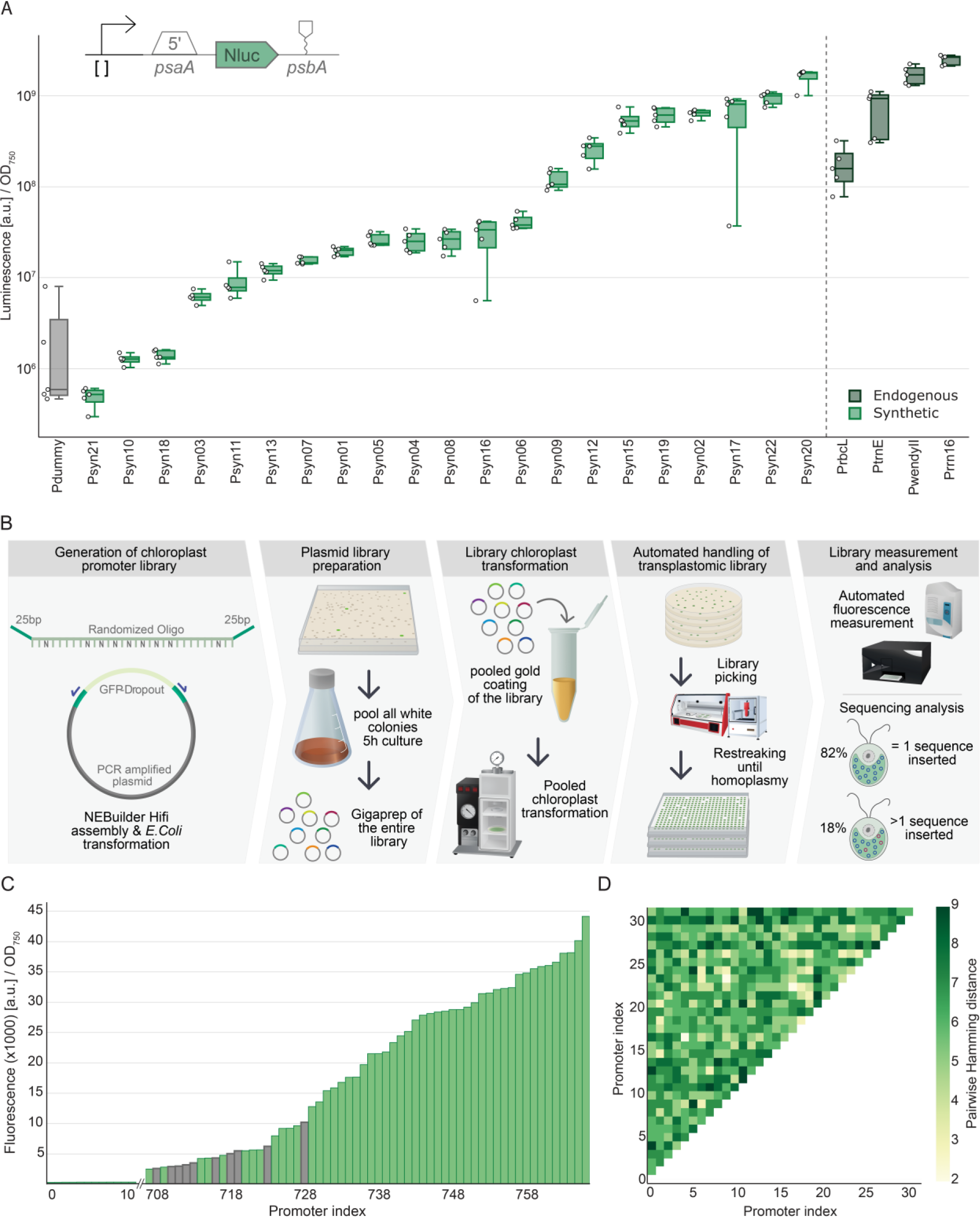
Designing and testing synthetic chloroplast promoters and library chloroplast transformation. **a,** Synthetic promoter characterization. Promoters were characterized by measuring NanoLuc activity of transplastomic strains containing different constructs, which differ in their promoters. Nanoluc signal of a transplastomic strain containing a NanoLuc expression cassette (green) is plotted as arbitrary units [a.u.] normalized to OD and compared to dummy sequence (gray). The dotted line demarcates synthetic promoters (left, light green) from endogenous ones (right, dark green). For all measurements, n_biological_= 5, n_technical_= 3. **b,** Conceptual workflow for transforming construct libraries into the chloroplast. Sequencing analysis showed that 82 % of transformants contained a single promoter variant and for 18% multiple sequence variants could be detected **c,** Fluorescence measurement of transplastomic promoter library. mScarlet-I fluorescence of the transplastomic strains containing different promoter variants was measured (Excitation 569 nm /Emission 605 nm). Fluorescence is plotted as arbitrary units [a.u.] normalized to OD _750_. 768 transplastomic strains have been measured and a range of different mScarlet-I signals can be detected. Gray bars represent examples that were selected in panel d for sequencing. **d**, Sequence analysis of sequenced transplastomic strains. Pairwise Hamming distance is plotted for the promoter sequence variant, which was found in a set of 33 sequenced transplastomic strains.

Even though above experiments successfully identified synthetic promoters, we note that the biolistic transformation remained a bottleneck in our automated workflow. This is, because each of the 21 promoters that we tested required an individual transformation, limiting the total number of promoter designs that could be assessed. To address this constraint, we explored the feasibility of a pooled library approach, as opposed to transforming individual constructs. As a proof of concept, we sought to transform and screen a library of degenerated promoter sequences. This library was created by cloning a degenerated single strand oligonucleotide into a pre-assembled vector containing a spectinomycin selection marker and an mScarlet expression cassette (Fig. 4b). After the cloning step, colonies were pooled, the DNA of the entire plasmid library was prepped and used for biolistic transformation. We selected 768 transformants that we systematically organized and screened via a fluorescent read-out. While 90% of the strains did show only little or no fluorescence, 10% showed a broad spectrum of mScarlet fluorescence. We continued to characterize selected transformant strains in more depth (Fig. 4c).

For that, we selected 32 individual strains and subjected them to sequencing. Using the individual promoter sequences of the library as barcode enabled us to analyse, whether individual sequence variants had been inserted or if a mixture of sequences persisted across the genome copies. Of these 32 strains, 26 strains (82%) contained one unique sequence, while the remaining six (18%) exhibited a mixture of sequences (Fig. 4d). Overall, these findings did not only provide more than 20 additional synthetic promoter sequences for the chloroplast, but notably also demonstrated the power of pooled library transformation in chloroplasts, opening new possibilities for plastid synthetic biology and engineering.

### Prototyping the effect of different genetic contexts on transgene expression

In all prior experiments, the different genetic elements were always assessed in the same genetic context, that is within the same transcription unit (and at one specific integration site), varying only one parameter (e.g., either promoter, 5’UTR, or 3’UTR). While this setup was helpful to study the individual behavior of the respective elements that were tested, for more advanced engineering approaches a collection of orthogonal part combinations are required that are unique in their respective sequence and can be utilized together in multi-transgene expression without the risk of unintended homologous recombination.

To prototype such orthogonal part combinations, we devised 11 unique combinations, all expressing the NanoLuc reported, but each varying in promoter, 5’UTR, and 3’UTR (Fig. 5a).

**Figure 5.**
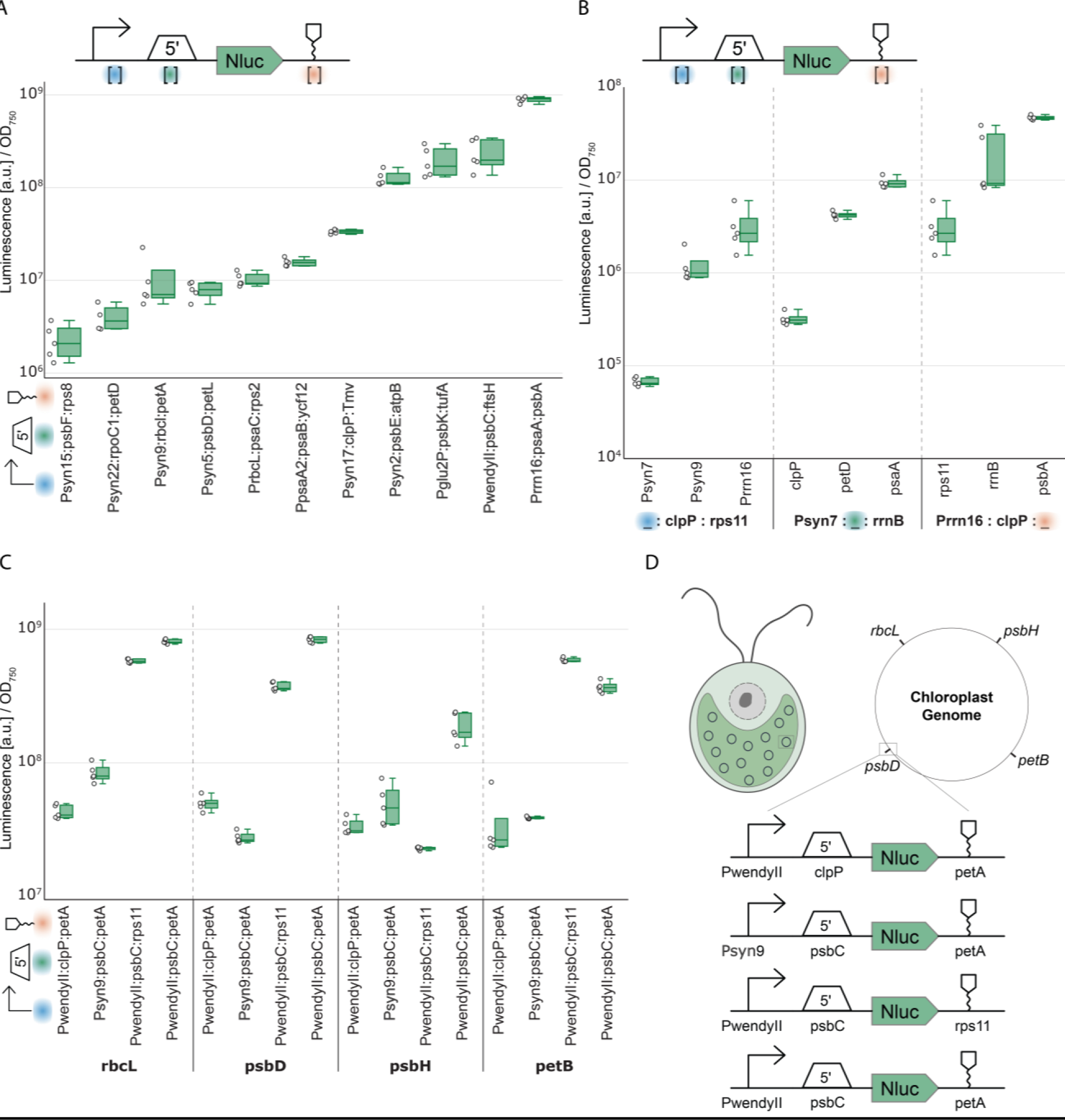
Characterizing the effects of different genetic contexts. **a,** Characterizing various combinations of parts. Combinations were characterized by measuring NanoLuc activity of transplastomic strains containing different constructs, which differ in their promoters, 5’UTRs and 3’UTRs. For clarity across the figure, part combinations are denoted as follows: Promoters by blue dots, 5’UTRs by green dots, and 3’UTRs by orange dots. Nanoluc signal of a transplastomic strain containing a NanoLuc expression cassette (green) is plotted as arbitrary units [a.u.] normalized to OD. For all measurements n_biological_= 5, n_technical_= 3. A range of three orders of magnitude of NLuc signal can be observed. **b,** Characterizing parts in weak expression backgrounds. Combinations were characterized by measuring NanoLuc activity of transplastomic strains containing different constructs, which differ in their promoters, 5’UTRs and 3’UTRs. Nanoluc signal of a transplastomic strain containing a NanoLuc expression cassette (green) is plotted as arbitrary units [a.u.] normalized to OD. For all measurements n_biological_= 5, n_technical_= 3. A range of two orders of magnitude of NLuc signal can be observed. **c,** Characterizing the effect of different sites of integration across the chloroplast genome integration sites were characterized by measuring NanoLuc activity of transplastomic strains containing different constructs, containing 4 different combinations of parts integrated in 4 different locations in the chloroplast genome. **d,** Conceptual design of the characterization of different integration sites. Four different constructs, which vary in the promoter, 5’UTR and 3’UTR are integrated in four different locations of the chloroplast genome.

In these experiments, the luminescence of the different combinations spanned three orders of magnitude, indicating a broad range of expression strength. In almost all cases, expression strength was correlated with expected 5’UTR activity (and promoter activity), with exception of the *SynP9:rbcL:petA* and *SynP17:clpP:ycf12* constructs, whose expression strength seemed to be rather correlated with expected promoter activity.

To study the context-dependency of individual genetic elements in more detail, we next devised a series of combinatorial constructs. This series included constructs, in which three different promoters of low-, medium-, and high-activity were tested in combination with low-activity elements clpP (5’UTR) and rps11 (3’UTR); three different 5’UTRs of low-, medium-, and high-activity were tested in context with the low-level activity promoter Psyn7 and the low-level activity 3’UTR rps11, respectively; and three different 3’UTRs of low-, medium-, high-activity were tested together with the high-activity promoter Prrn16 and the low-activity 5’UTR clpP. In all cases, the elements behaved as expected by increasing expression strength by approximately 10-fold and 50-fold for the medium- and high-activity element, confirming their functionality across different genetic contexts (Fig. 5b).

Finally, we also studied the impact of different integration sites within the chloroplast genome. To this end, we created four different transcriptional units (two of low and two of high transgene expression strength) that we integrated into four distinct locations (Fig. 5d, (Extended Data Fig. 4) within the chloroplast genome (psbH, rbcL, psbD, petB). For the rbcL, psbD, and petB integration sites we observed comparable expression levels between the four constructs (Fig. 5c). However, for the psbH integration site, we noted some differences, particularly for the construct containing the rps11 3’UTR. This observation is in line with a recent report that the 3’UTR at the psbH site is important in controlling read-through activity of neighboring genes, which in turn can modulate transgene expression.^67^ Overall, these experiments showed that most constructs behaved similar independent of their genomic position, but also highlighted that validating individual constructs at the psbH site is important for quantifying actual transgene expression strength.

### Applying the tools to prototype novel chloroplast traits in *C. reinhardtii*

Finally, we used our plastome engineering toolkit to demonstrate the prototyping of novel metabolic traits in the chloroplast of *C. reinhardtii*. As case example, we sought of introducing a recently reported synthetic photorespiratory bypass^7^ into the chloroplast that is based on two enzymes: glycolate dehydrogenase (GDH) and malate synthase (MS), respectively (Fig. 6a). These two enzymes cause the direct decarboxylation of the photorespiration product glycolate into CO_2_, which increases CO_2_ concentrations in the chloroplast and was reported to improve photosynthetic yield in different crops.^7,68^

**Figure 6.**
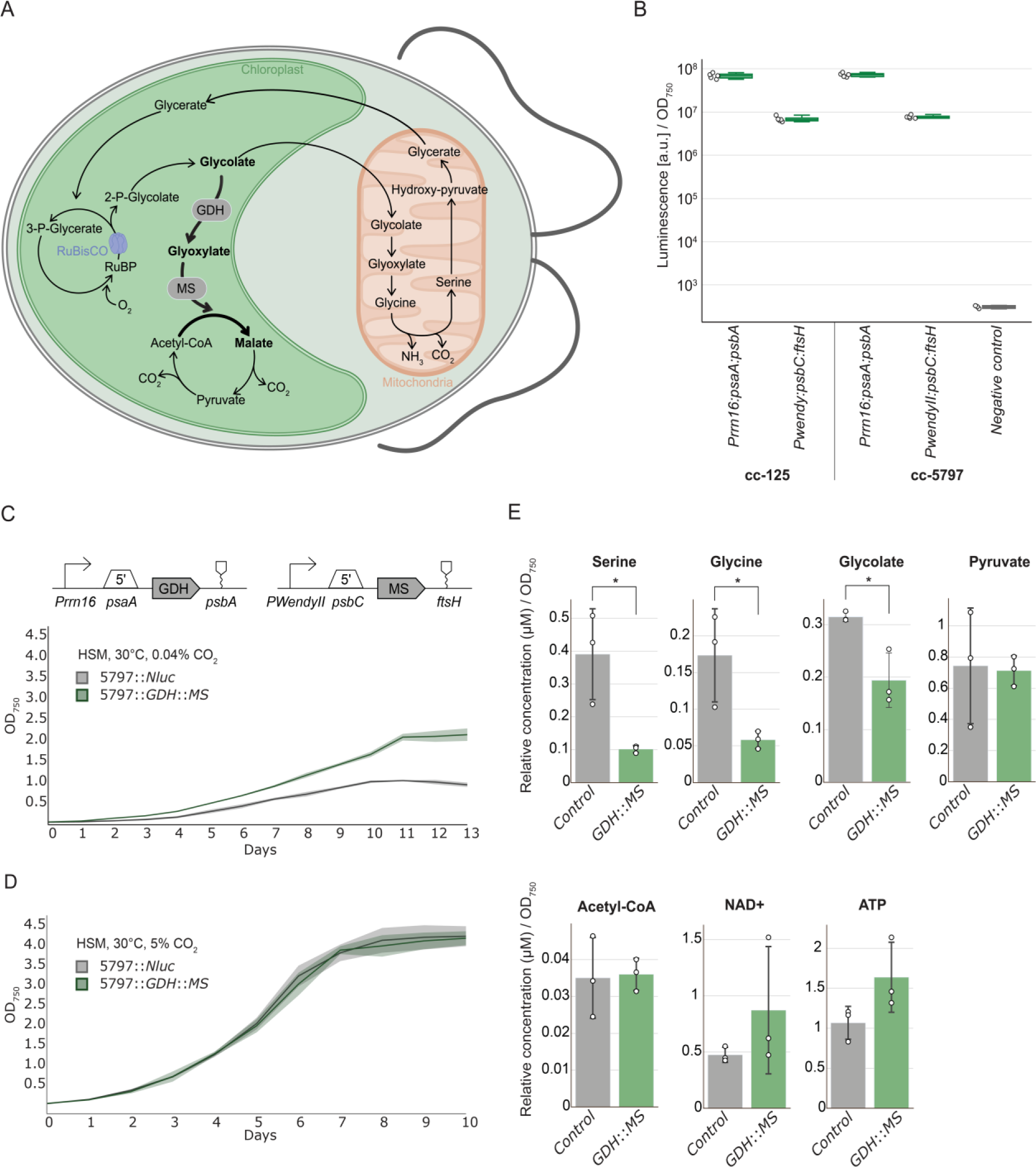
Introducing a synthetic photorespiratory bypass in the chloroplast of *C. reinhardtii*. **a**, Metabolic scheme for the previously reported synthetic photorespiration bypass. The pathway relies on the introduction of two genes (in gray): the glycolate dehydrogenase (GDH) and malate synthase (MS). **b,** Nanoluc assay for part combinations characterization in the cc-125 WT and cc-5797 mutant. Nanoluc signal is plotted as arbitrary units [a.u] normalized to OD_750_ and compared to the control-lacking reporter (in gray). For all measurements, n_biological_=5. **c-d**, Growth curve of engineered strains and control at ambient CO2 and 5% CO2 respectively, shown as OD_750_ over time (days). Parts tested in b were used to express both GDH and MS. Engineered strains (5797::GDH::MS,in green) reach a two-fold final OD over the control strains (5797::Nluc) in ambient CO2 but behave similarly in 5% CO2. n_biological_=3 for each curve. **e.** Intracellular metabolome analysis of engineered and control strains during the exponential growth phase. Relative concentrations (μM/OD_750_) of serine, glycine, glycolate, pyruvate, acetyl-CoA, NAD+ and ATP are displayed for the 5797::GDH::MS (in green) and 5797::Nluc (in gray). n_biological_=3 for each experiment. Significant changes are indicated by a * for a p_value<0.05. (serine, p_value = 0.023. glycine, p_value = 0.037. glycolate, p_value = 0.017).

As host for our engineering efforts, we chose a CO_2_-concentrating mechanism deficient- and glycolate dehydrogenase-deficient *C. reinhardtii* mutant (cc-5797, genotype confirmation in Supplementary Fig. 2c-d) that shows a severe growth phenotype under atmospheric CO_2_ concentrations due to increased photorespiration^69^. To express GDH and MS from the chloroplast genome, we chose four regulatory elements and validated their activity at the psbH integration site in cc-5797 using the NanoLuc reporter (Fig. 6b). We assembled the transcriptional units for expression of GDH and MS, integrated these units into the psbH locus, confirmed integration via colony PCR (Extended Data Fig. 5a) and verified the presence of the malate synthase and the glycolate dehydrogenase through proteomics ((Extended Data Fig. 5c). Interestingly, during our analysis, we detected functional expression of glycolate dehydrogenase, indicating that the 5’UTR knock out of glycolate dehydrogenase was not completely tight in the cc 5797 strain, suggesting that the growth phenotype was mainly caused by deficiency of the CO_2_-concentrating mechanism.

At 5% CO_2_ concentration, the *C. reinhardtii* strain expressing MS and GDH and the negative control (i.e., the cc 5797 strain expressing NanoLuc luciferase) behaved identical (Fig. 6d). However, at ambient CO_2_ concentrations, the strain with the functional photorespiration bypass showed a clear growth benefit (Fig. 6c, Extended Data Fig. 5b). While both strains had comparable doubling times (∼43 hours), the strain expressing MS and GDH reached almost two-fold higher final ODs, indicating that the pathway was functionally expressed from the chloroplast (Fig. 6c, (Extended Data Fig5.b). Moreover, intracellular metabolomics unraveled distinct differences between the engineered and the control strain. In the engineered strain, intracellular glycolate concentrations (as well as the derived amino acids glycine and serine) were significantly decreased compared to the control strain (Fig. 6e). In contrast, the levels of pyruvate and energy metabolites, such as ATP were not significantly different between the two strains (Fig. 6e), supporting the hypothesis that expression of MS and GDH had re-channeled photorespiration flux towards the synthetic glycolate metabolism. Together, these results demonstrated the general possibility to prototype novel metabolic traits, such as improved photorespiration, in the chloroplast of *C. reinhardtii*.

## Discussion

Harnessing the full potential of synthetic biology in photosynthetic eukaryotes (and plant systems in particular) necessitates considerable improvements in genetic engineering capabilities, with chloroplasts emerging as a prime target for such efforts. Traditionally, chloroplast engineering has been limited to a relatively small number of transplastomic strains or plant lines, which has challenged a systematic characterization of genetic parts and more complex constructs in chloroplasts.^32^

In this study we established *C. reinhardtii* as chloroplast model system, implemented an automated workflow for the generation and handling of thousands transplastomic strains in parallel, and introduced an extensive array of genetic tools to significantly advance the field of chloroplast synthetic biology. By dramatically expanding the number of selection of markers, reporter genes, and regulatory elements, our study paves the way for more ambitious chloroplast engineering projects in *C. reinhardtii*. To facilitate a broader application and further development within the community, all tools, including the 300 genetic parts are available through repositories, and are already in use by other laboratories.^27^ This set of parts provides a first reference set that can be systematically extended in the future by more high-throughput screens, including pooled library approaches, which have now become accessible for chloroplasts through our efforts.

Our comprehensive toolkit enables multi-transgene expression through verified part combinations, and offers the possibility for more advanced transgene stacking in the plastid of *C. reinhardtii*. Thus, chloroplast-based strategies become a viable alternative for applications targeting the chloroplast compared to nuclear genome-based efforts, which have been used so far, but suffer from several disadvantages, such as low transgene expression and gene silencing.^70,71^ We demonstrate such an application by establishing a recently reported synthetic photorespiration pathway^7^ in *C. reinhardtii* chloroplasts that leads to a twofold increase biomass yield under selection. This proof-of-concept provides new possibilities for ongoing efforts to establish novel synthetic photorespiration pathways in chloroplasts (e.g., the TaCo pathway^5^ or BHAC cycle^8,9^, or even replacing the Calvin cycle by new-to-nature carbon fixation pathways (e.g., the Theta cycle^12^ or reverse glyoxylate shunt.^72^

Despite some differences between the chloroplast of *C. reinhardtii* and land plants, the plastome structure shows a high degree of conservation across photosynthetic eukaryotes. Genetic elements from the chloroplast of *C. reinhardtii* have been successfully used in various land plants, including tobacco^20^, *Arabidopsis*^45^, and poplar^43,44^, and *vice versa* (e.g., as shown in this study). This high degree of transferability lays the basis for the rapid testing and optimization of genetic designs through our prototyping system, before a transfer of results to land plants and crops. Especially when using high-throughput routines combined with pooled library strategies, these efforts could dramatically expand the search for new phenotypes and improved chloroplast functions, which could significantly contribute to developing climate-resilient crops, addressing agricultural challenges for an ever-growing world population facing climate change.

## Materials and Methods

### *Escherichia coli* Strains, Transformation, and Growth Conditions

Bacterial cultures during the cloning phase were incubated at 37°C in Luria-Bertani (LB) broth under shaking conditions for liquid cultures and on LB agar plates with 1.5% (m/V) agar for solid cultures. Antibiotic selection was applied according to the construction level: Chloramphenicol (34 µg/mL) for level 0 constructs (pME_Cp_UAV_GFP), Ampicillin (100 µg/mL) for level 1 (pME_Cp_0_7-8_003), and Kanamycin (50 µg/mL) for level 2 (pME_CP_0_7-8_006). Two chemically competent *E. coli* strains, DH10B and NEB Turbo, both sourced from New England Biolabs, were utilized for transformations, which were performed via heat shock as per the manufacturer’s instructions. Plasmid DNA was isolated using Macherey-Nagel kits designed for both mini and midi preparations, the latter being essential for chloroplast transformation due to the supercoiled nature of the DNA obtained. Terrific Broth (TB) was employed to maximize yields in midi preparations.

### *Chlamydomonas reinhardtii* Strains and Growth Conditions

Wild-type strains of *Chlamydomonas* were cultured on Tris-Acetate-Phosphate (TAP) medium in high light (∼100 μmol photons m^-2^ s^-1^), following the Chlamydomonas Resource Center’s recipe, which included Hutneŕs trace elements^73^. The medium was supplemented with Ampicillin (100-500 µg/mL) and Difco agar (Beckton Dickson, 0.8% m/V) for solid cultures. Transformations were performed on TAP agar plates supplemented with spectinomycin (100 µg/mL), tobramycin (100 µg/mL), or kanamycin (200 µg/mL) as required. Tobramycin antibiotic stocks have been prepared freshly every time. Transformed strains were propagated on TAP medium supplemented with Ampicillin (500 µg/mL) and an increasing concentration of either spectinomycin (100, 300, 500, 1000 µg/mL) or tobramycin (100, 150, 300 µg/mL) during the restreaking phase. Culture flasks were meticulously cleaned with detergent, rinsed thrice with deionized water, and autoclaved with Milli-Q water to eliminate any residual detergent (SOMAT). The study predominantly utilized the CC-125 mt+ mutant strain from the Chlamydomonas Resource Center, as well as cc-5797, a double mutant for Glycolate dehydrogenase and cia5, which is a global regulator for the CO_2_ concentrating mechanism in *C. reinhardtii*.

For growth curves of cc-5797, pre-cultures were grown for 9 days in High Salt Medium, according to the Chlamydomonas Resource Center (HSM + Spec_100_, 5% CO_2_, 30°C) and used for inoculation of cultures at OD_750_=0.05. Growth curves were carried in HSM supplemented with Spec_100_, 0.04% CO_2_, 30°C or HSM + Spec_100_, 5% CO_2_, 30°C for the control. 1 mL samples were taken every day and OD_750_ was measured.

### Chloroplast Transformation

Chloroplast transformation in *Chlamydomonas* was executed using a Bio-Rad PDS/1000 He (Bio-Rad) gene gun. Briefly, cells were grown in 50 mL TAP medium for two days before inoculation into two 400 mL TAP cultures with 10-15 mL of the pre-culture. These cultures were grown to a density of 10^7^ cells/mL, harvested by centrifugation at 3000x g for 10 minutes, and the pellets resuspended in 50 mL fresh TAP medium to disperse clumps before a second centrifugation at 3000x g for 10 minutes. Pellets were resuspended in 10 mL TAP medium, and 500 µL was used for plating on TAP agar plates supplemented with the appropriate selection antibiotics. Plates were dried, uncovered, for at least one hour. 60 mg of gold particles (0,6 µm Bio-Rad) was weighed and washed with 100µl 100% ethanol. Subsequently gold particles were washed twice (1mL of water) and resuspended in 1mL of water. 50µl of washed gold particles were coated with ∼10 μg of supercoiled plasmid DNA prepared via midi preparation for each construct, using 50µl of 2.5 M CaCl2 and 10µl 0.1 M spermidine. All reagents were kept on ice throughout the protocol prevent particle clumping. A pure ethanol wash was carried out, and DNA-bound gold beads were resuspended in 60µl pure ethanol and used for biolistics, following Bio-Rad instructions. 1100 psi rupture disks were used. Plates were incubated in the dark for recovery (∼5-10 μmol photons m^-2^ s^-1^) for 24h and moved to high light for selection (∼100 μmol photons m^-2^ s^-1^).

### Automated Picking, Re-streaking, and Re-arraying

A PIXL robot (Singer Instruments) facilitated semi-automatic colony picking across various array formats, including 96, 384, or 1536 configurations, with optimized parameters ensuring consistent selection of *Chlamydomonas* colonies as outlined in the Supplementary Text 2. For each construct, 16 colonies were selected when possible. The Rotor (Singer Instruments) played a crucial role in managing arrays of transformants, adjusting antibiotic concentrations, and preparing assay plates. It was employed for restreaking tasks and for dividing larger 384-well plates into smaller 96-well measurement arrays, utilizing pin formats compatible with both plate sizes.

### Part Design

For our **Chl**amydomonas **Chloro**plast **Mod**ular **A**ssembly **S**ystem (CHLOROMODAS), we employed the Phytobrick standard for the design of Level 0 parts^47^, as previously utilized in other plant and Chlamydomonas MoClo toolkits^37,46,74^. To assemble multiple transcription units, we adapted the standard of the Marburg collection^75^, with the exception of exchanging the 3’ Connector/Backbone Part overhang from AGCT to AGAC to enhance assembly efficiency as determined by the NEB ligase fidelity tool.^76^ All Level 0 parts were cloned into a standard universal acceptor vector.

For the rational design of 5’UTR and 3’UTR Level 0 Parts, we utilized previously reported RNA-seq and sRNA data^60^ from C. reinhardtii chloroplasts for the annotation of transcription start sites (TSS) and RNA-binding sites of trans-factors. Additionally, RNA secondary structures of 3’UTRs were analyzed using ViennaRNA Web Services to identify necessary stabilizing hairpin structures. The design of endogenous promoter parts involved examining RNA-seq data to identify consensus −35 and −10 promoter sequences in the plastome prior to the TSS. Suitable candidates for intercistronic expression elements (IEEs) were identified by analyzing co-transcription data from a list of genes in the chloroplast and searching for RNA-binding protein footprints within these sequences, indicating potential post-transcriptional processing success.

Several parts in our toolkit underwent multiple design-build-test cycles using reporter gene assays to determine the minimal sequence required for optimal function, such as for the rrn16 promoter and rbcL 3’UTR. Codon usage for coding sequences was optimized using the Geneious codon optimization tool based on the chloroplast codon usage table for C. reinhardtii (available at https://www.kazusa.or.jp). Original sequences for the glycolate dehydrogenase (GDH) and malate synthase (MS) were sourced from South et al., 2019 and subsequently optimized.

Level 0 parts were either amplified from the cc-125 genome or synthesized as fragments by Twist Bioscience. Smaller parts (<100 bp) were cloned using annealed oligonucleotides.

### Part assembly

Vectors and inserts were standardized to concentrations of 20 fmol. The assembly of Level 0 and Level 2 constructs utilized the type IIS restriction enzyme BsmBI, while BsaI was employed for the assembly of Level 1 constructs. These assembly reactions were facilitated using the NEBridge Golden Gate Assembly Kit from New England Biolabs for both BsaI and BsmBI enzymes. The reaction mixtures, prepared without specifying volumes, included a vector at 10 fmol, inserts at 20 fmol, the respective NEBridge Golden Gate Enzyme Mix (BsaI or BsmBI), T4 DNA Ligase Reaction Buffer, and nuclease-free water to complete the mixture. BsaI-based assembly reactions underwent 50 cycles of temperature alternations, with periods at 37°C followed by cooling to 16°C. In contrast, BsmBI-based reactions alternated between 42°C and 16°C. The final steps for both methods involved a brief incubation at 60°C and a final heat inactivation at 80°C before the transformation into competent bacterial cells (5µl of the Golden Gate mixture).

### Fluorescence assays & FACS

For chloroplast fluorescence analysis in *Chlamydomonas reinhardtii*, transformed and wild-type (WT) colonies from 384-well arrays were resuspended in nuclease-free water in black 384-well Greiner Bio-One plates. Optical density (OD) was measured at 750 nm for normalization. Fluorescence quantification utilized a microplate reader (Tecan infinite 200 Pro) with specific settings for mScarlet-I (excitation: 569 nm, emission: 605 nm), mVenus (excitation: 515 nm, emission: 545 nm), and tdTomato (excitation: 554 nm, emission: 585 nm).

FACS analysis was performed using a SONY SH800S Cell Sorter to assess fluorescent protein expression. WT and mScarlet-I expressing strains underwent a two-day pre-culture and a two-day culture in TAP medium, with spectinomycin (100 µg/mL) added for transformants. Samples were diluted tenfold in TAP for optimal sorting. The SONY SH800S settings were adjusted to accurately detect the fluorescent markers according to manufacturer’s instructions, which can be found in the Supplementary text 2.

### High-throughput pipeline for luminescence measurements

To facilitate the high-throughput analysis of regulatory elements, a detailed pipeline was established, as illustrated in Figure 1. This approach incorporated NanoLuc luciferase assays from Promega for the quantitative assessment of regulatory element functionality. Initially, transformants were propagated on 384-well agar plates. These plates were then segmented into four 96-well solid-agar arrays with the aid of the Rotor robot, enhancing the biomass by replicating each transformant into 5×5 squares. This cultivation extended over an 8-day period to ensure adequate growth. Post-cultivation, colonies were carefully resuspended in 30 µL of water within 384-well ECHO 525 plates (LABCYTE). Optical density (OD) measurements at 750 nm were conducted to ascertain cell density. To ensure standardized measurements across seven replicates per construct, a custom Python pipeline was created. This pipeline leverages a configuration file to set the liquid handling parameters for the ECHO 525 and define experimental variables, including number of technical and biological replicates. The objective was to achieve an OD_750_ of 0.01 for each replicate. To this end, the script uses the previously measured absorbance data to generate pipetting instructions for the ECHO 525 liquid handler, which include automated modifications to the plate layout and the selection of source wells. The dilutions were created in white 384-well plates, with alternate wells left empty to minimize light interference between replicates in the subsequent luminescence measurements. The luminescence assay was initiated by adding 10 µL of Nanoluc substrate solution to each well. Following a 5-minute incubation at room temperature, luminescence was measured using a plate reader. Afterwards, the data processing pipeline utilized a JSON mapping file to link each measurement with the corresponding construct. By integrating the ECHO pipetting instructions with the absorbance and luminescence data, a comprehensive experiment overview was automatically compiled. This document includes normalized luminescence values for each construct, corrected based on the OD_750_ measurements and dilution factors to precisely reflect the activity of regulatory elements. This overview was subsequently used for further analysis and data visualization.

### Metabolome extraction

Daily 1 mL samples from the growth curve were mixed with an equal volume of 70% methanol pre-cooled at −70°C, centrifuged, and the supernatant discarded before freezing the pellets at −70°C. In a chemical fume hood and with proper PPE (gloves and glasses), 200 µL of 50% methanol in TE buffer (10 mM Tris-HCl, 1 mM EDTA, pH 7.0) was added to each pellet. Mechanical disruption was performed in a bead mill (Bead Bug) using 0.375 g of acid-washed glass beads (500 µm) for two 30-second cycles at maximum power, with samples cooled on ice between cycles. Following disruption, 200 µL of 100% chloroform was added, and the mixture was vortexed for 30 seconds. After centrifugation at 13,000 xg for 10 minutes at −10°C, the upper phase was extracted with 1 mL syringes, slowly filtered through PTFE filters into pre-cooled Eppendorf tubes, and transferred into metabolomics vials for analysis, with careful handling to avoid warming and contamination.

### Metabolome analysis

Quantitative determination of the targets was performed using an LC-MS/MS system, integrating chromatographic separation and mass spectrometry for precise analysis. Chromatographic separation was achieved on an Agilent Infinity II 1290 HPLC system, employing columns of varying specificities (Kinetex EVO C18, ZicHILIC SeQuant, and SeQuant ZIC-pHILIC) with particle sizes ranging from 3 to 5 μm and pore sizes of 100 Å (Phenomenex), connected to guard columns of similar specificity. The flow rates varied between 0.1 and 0.3 mL/min, with mobile phases tailored to each experiment: CoA esters (1) 50 mM Ammonium Formate in water and 100% methanol, organic acids (2) 0.1% formic acid in water and methanol, amino acids (3) 0.1% Formic acid in 99:1 water:acetonitrile and acetonitrile:water, and ATP and NAD+ (4) 10 mM ammonium acetate in water and acetonitrile, all adjusted to specific pH levels and temperatures. Injection volumes were set at 1 or 2 µL, with mobile phase profiles designed to optimize separation over varying times and gradients.

Mass spectrometry analysis was conducted using an Agilent 6495 ion funnel mass spectrometer, operating in both positive and negative ionization modes with specific settings for ESI spray voltage, nozzle voltage, sheath gas temperature, nebulizer pressure, and drying gas temperature to ensure optimal detection and quantification. Compounds were identified and quantified based on their mass transition, retention time, and peak area, compared to standards, using MassHunter software. Relative abundance and absolute concentrations were determined based on peak areas and external standard curves, respectively. The optimization of mass transitions, collision energies, cell accelerator voltages, and dwell times was achieved using chemically pure standards, with the parameter settings detailed in Supplementary Tables 1 through 4 for each experimental variation.

### Proteome extraction

Cells were cultured in TAP medium for four days until reaching a density of 10^7^ cells/mL or exponential phase in a 50 mL pre-culture. Cultures was centrifuged at 3000x g for 10 minutes at room temperature. The supernatant was removed, and the cell pellet was washed with 30 mM Tris-HCl (pH 7.9), followed by a second centrifugation. The pellet was then either used immediately or stored at −80°C. Cell pellets were lysed by adding 500 µL of Tris-HCl with protease inhibitors and 0.5 µg/µL lysozyme (Sigma-Aldrich), incubating at 25°C for 1 hour. Then, 400 µL of 2% sodium deoxycholate (SDC) with 10 mM dithiothreitol (DTT) was added. After three freeze-thaw cycles, the pellet was sonicated, heated at 90°C for 30-45 minutes, and sonicated again. The lysate was centrifuged at 14,000x g for 30 minutes at 4°C, collecting the supernatant for further processing. The supernatant was mixed with six volumes of cold (−20°C) acetone and incubated overnight at −20°C. After centrifugation at 5000x g for 20 minutes at 4°C, the supernatant was discarded. The pellet was washed twice with cold methanol, dried, and reconstituted in 200 µL of 0.5% SDC. Protein concentration was determined using BCA and pre-diluted BSA (Pierce BCA, Thermo Fischer Scientific) following manufacturer’s protocol, and 50 µg total protein was used for further protein purification by SP3^77^ as described previously with minor modifications. In short, proteins were bound to 4 µl SP3 beads (40% v/v) in presence of 70 % acetonitrile for 15 min at room temperature, followed by two washes of beads with 70 % Ethanol and an additional wash with acetonitrile. After removal of the supernatant, 1 µg trypsin in 100 mM NH_4_HCO_3_ were added to the beads and digested shaking overnight at 30°C. Following digest, beads were separated and peptides containing supernatant was collected and further purified and desalted by C18-solid phase extraction using Chromabound spin columns (Macherey-Nagel). Cartridges were prepared by adding acetonitrile, followed by equilibration with 0.1% TFA. Acidified peptides were loaded on equilibrated cartridges, washed with 5% ACN and 0.1% TFA containing buffer and finally eluted with 50% ACN and 0.1% TFA.

### Proteome analysis

Dried peptides were reconstituted in 0.1% Trifluoroacetic acid and then analyzed using liquid-chromatography-mass spectrometry carried out on an Exploris 480 instrument connected to an Ultimate 3000 RSLC nano and a nanospray flex ion source (all Thermo Scientific). Peptide separation was performed on a reverse phase HPLC column (75 µm x 42 cm) packed in-house with C18 resin (2.4 µm; Dr. Maisch). The following separating gradient was used: 94% solvent A (0.15% formic acid) and 6% solvent B (99.85% acetonitrile, 0.15% formic acid) to 25% solvent B over 97 min., and an additional increase of solvent B to 35% for 35 min. at a flow rate of 300 nl/min. MS raw data was acquired on an Exploris 480 (Thermo Scientific) in data independent acquisition (DIA) mode. Peptides were ionized at a spray voltage of 2.3 kV, ion transfer tube temperature set at 275 °C. The funnel RF level was set to 45. For DIA experiments full MS resolutions were set to 120.000 at m/z 200 and full MS, AGC (Automatic Gain Control) target was 300% with an IT of 50 ms. Mass range was set to 350-1200. AGC target value for fragment spectra was set at 3000%. 60 windows of 4 Da were used with an overlap of 1 Da (m/z range 550-678). Resolution was set to 15,000 and IT to 50 ms. Stepped HCD collision energy of 27, 30, 32 % was used. MS1 data was acquired in profile, MS2 DIA data in centroid mode. Analysis of DIA data was performed using the DIA-NN version 1.8^78^ a uniprot protein database from *Chlamydomonas reinhardtii* with additionally added sequences to generate a data set specific spectral library for the DIA analysis. The neural network based DIA-NN suite performed noise interference correction (mass correction, RT prediction and precursor/fragment co-elution correlation) and peptide precursor signal extraction of the DIA-NN raw data. The following parameters were used: Full tryptic digest was allowed with two missed cleavage sites, and oxidized methionines and carbamidomethylated cysteins. Match between runs and remove likely interferences were enabled. The precursor FDR was set to 1%. The neural network classifier was set to the single-pass mode, and protein inference was based on genes. Quantification strategy was set to any LC (high accuracy). Cross-run normalization was set to RT-dependent. Library generation was set to smart profiling. DIA-NN outputs were further evaluated using the SafeQuant^79,80^ script modified to process DIA-NN outputs

### Immunoprecipitation analysis

For protein analysis via SDS-PAGE, samples were prepared by centrifuging 8 ml of culture with 4.10^7^ cells at 4500 rpm and 4°C for 2 minutes, decanting the supernatant, and resuspending the cells in the remaining medium for transfer to 1.5 ml Eppendorf tubes. After a subsequent centrifugation for 30 seconds at 4°C, the supernatant was removed, and cells were resuspended in 120 µl of 0.1 M Na2CO3 carbonate buffer, followed by the addition of 110 µl of 5% SDS solution. Samples were then mixed vigorously, heated at 95°C for 45 seconds, snap-frozen in liquid nitrogen, and stored at −20°C.

Protein concentrations were determined using the Lowry assay. Solutions A and B for the Lowry assay were prepared and mixed, stored at 4°C. Protein standards were prepared by diluting 1 mg/ml BSA in ddH2O, and samples were prepared from the −20°C freezer, with triplicates of 90 µl water and 10 µl supernatant. After adding 1 ml of Lowry’s solution and incubating for 15 minutes at 25°C, 100 µl of 1x Folin reagent was added, and samples were incubated for an additional 20 minutes. Absorbance was measured at λ = 750 nm to determine protein concentrations.

SDS-gel electrophoresis was performed using prepared solutions for stacking (3%) and separating (10%) gels. Gels were assembled, polymerized, and loaded with samples prepared with 30 µg of protein in a final volume of 60 µl, including 4x loading buffer. Electrophoresis was conducted at 160 V until the chlorophyll was about to exit the gel. Subsequently, samples were prepared for Western blotting using Whatman paper and nitrocellulose membranes. The membranes were blocked with 5% non-fat dry milk in TBST (20 mM Tris, 150 mM NaCl, 0.1% Tween-20) for 1 hour at room temperature and incubated overnight at 4°C with primary antibodies specific to the target tag: HA tag, mouse monoclonal antibody (Sigma). FLAG, mouse monoclonal antibody (Sigma). Myc, mouse monoclonal antibody (Invitrogen). Following primary antibody incubation, membranes were washed three times with TBST and secondary antibodies for 1 hour at room temperature and proteins were analyzed using a gel imager.

### Data analysis, plotting and statistical analysis

Data analysis and visualization were performed using Python 3.10.5. For parsing and processing, the pandas library (version 1.4.3) was utilized. Visualization of the data was conducted using the Plotly library (version 5.9.0). Data is displayed as box plots and adjacent individual data points on decadic logarithm scale. The midlines of the box plots represent the median, the boxes’ upper and lower limits represent the 1st and 3rd quartiles. Whiskers correspond to the box’ edges +/-1.5 times the interquartile range.

Bar graphs, specifically in Figures 2 and 6, depicted mean values with error bars indicating standard deviation, providing a clear visual summary of the data’s central tendency and variability. For the analysis of fluorescence-activated cell sorting (FACS) data, the FlowCal library (version 1.3.0) was employed, presenting data on a logicle scale, a method detailed in the provided reference.^81^

Metabolomics data underwent statistical examination using the scipy library (version 1.8.1). The significance of these analyses was based on unpaired t-tests. Growth curve data were visualized using mean values with shading to represent the standard error. Doubling times for each strain were calculated using GraphPad Prism 10 and its Exponential (Malthusian) Growth function, providing the doubling time over the first 7 days (exponential phase).

### Fluorescence microscopy

For imaging, cells were grown to exponential phase (2-8×10^6^ cells mL^-1^) in TAP media. 10 μL of cell suspension was overlaid with 30 μL of 1% TP-low-melting point agarose in µ-Slide 18 Well chambered coverslips (ibidi, Germany). All fluorescence images were captured using a Zeiss LSM880 confocal microscope in Airyscan mode using a 63x 1.4 numerical aperture (NA) Plan-Apo oil-immersion lens (Carl Zeiss, Germany). Brightfield images were captured using the microscope T-PMT output channel. Specific settings are given in Supplementary text 1.

### Colony PCR analysis

Chlamydomonas strains were subjected to colony PCR analysis. The cells were processed in 384 Greiner PCR plates, each well containing 25 μL of an extraction buffer composed of 10 mM Tris, 5 mM EDTA, and 0.01% Triton100, adjusted to pH 8. The plates underwent vortexing, followed by a boiling step at 100°C for 10 minutes. Subsequently, the plates were vortexed again and then cooled on ice for 5 minutes before centrifugation. The supernatant obtained from this process was diluted in a 1:5 ratio with the extraction buffer and used as the DNA template for PCR reactions.

For the PCR amplification, the 384 Greiner PCR plates were prepared by adding 0,5μL of primer (10 μM) to each well, followed by 8 μL of Phusion High Fidelity Master mix (New England Biolabs) according to the manufacturer’s instructions. These steps were facilitated using the ECHO 525 liquid handling system. To this mixture, 1 μL of the 1:5 diluted DNA template was added. The PCR reaction was then performed as per the manufacturer’s guidelines, with the annealing temperature being adjusted based on the specific primer combination used. GeneRuler 1kb DNA ladder (Thermo Fischer Scientific) was used for every gel shown in the study. cPCR confirming Nanoluc insertion in each strain used for part characterization can be found in Supplementary Fig1 and 2.

### Design, construction, and chloroplast transformation of a degenerated promoter library

A 90 bp single stranded degenerated oligo was designed and ordered. This oligo has 25bp homologies at the 5’ and 3’ end to an already pre-assembled vector. The oligo also contains 40 bp of the core *rrn16* promoter, but with 9 sites which are changed to a degenerated base pair. The pre-assembled vector contains a spectinomycin resistance and a mScarlet-I expression cassette, consisting of a *psaA* 5’UTR, the mScarlet-I coding sequence, the *psbA* 3’UTR and 5’and 3’ homology flanks for the integration in the *psbH* locus. The vector was then PCR amplified and subsequently used for the NEB HiFi assembly together with the degenerated oligo. Afterwards all red colonies were pooled in 500mL of TB medium. After 5 hours of incubation, the DNA of the entire library was isolated via the NucleoBond Xtra Maxi Kit (Macherey Nagel). This pooled DNA-isolation was used for the subsequent chloroplast by coating the plasmid mixture on the gold nanoparticles. After the chloroplast transformation, 768 colonies were picked and restreaked 4 times (once a week) to reach homoplasmy.

## Data availability

Raw mass spectrometry data, as well as annotated plasmid maps will be submitted to the “Edmund” Max Planck Society open-source database, as well as deposited in GenBank.

## Code availability

The automation and data analysis scripts as well as the genbank files of all the plasmids used in this study (including the entire chloroplast MoClo collection) are available at https://github.com/ChlamyMarburg/ChloroplastTools

## Acknowledgements

We thank Vinca Seiler for creating all the illustrations of the manuscript.

## Author information

These authors contributed equally: René Inckemann and Tanguy Chotel

## Authors and Affiliations

**René Inckemann** - Max-Planck Institute for Terrestrial Microbiology, Marburg 35043, Germany; Center for Synthetic Microbiology, Philipps-Universität Marburg, Marburg 35032, Germany

**Tanguy Chotel** - Max-Planck Institute for Terrestrial Microbiology, Marburg 35043, Germany; Center for Synthetic Microbiology, Philipps-Universität Marburg, Marburg 35032, Germany

**Cedric K. Brinkmann -** Max-Planck Institute for Terrestrial Microbiology, Marburg 35043 Germany;

**Michael Burgis** - Center for Synthetic Microbiology, Philipps-Universität Marburg, Marburg 35032, Germany;

**Laura Andreas -** Max-Planck Institute for Terrestrial Microbiology, Marburg 35043, Philipps-Universität Marburg, Marburg 35032, Germany;

**Jessica Baumann -** Max-Planck Institute for Terrestrial Microbiology, Marburg 35043, Philipps-Universität Marburg, Marburg 35032, Germany;

**Priyati Sharma -** Max-Planck Institute for Terrestrial Microbiology, Marburg 35043

**Melanie Klose -** Max-Planck Institute for Terrestrial Microbiology, Marburg 35043, Germany;

**James Barret -** Centre for Novel Agricultural Products (CNAP), University of York, York YO10 5DD

**Fabian Ries-** TU Kaiserslautern, Kaiserslautern 67663, Germany

**Nicole Paczia -** Max-Planck Institute for Terrestrial Microbiology, Marburg 35043, Germany

**Timo Glatter -** Max-Planck Institute for Terrestrial Microbiology, Marburg 35043, Germany

**Luke Mackinder -** Centre for Novel Agricultural Products (CNAP), University of York, York YO10 5DD

**Felix Willmund -** Philipps-Universität Marburg, Marburg 35032, Germany; TU Kaiserslautern, Kaiserslautern 67663, Germany

**Tobias J. Erb** - Max-Planck Institute for Terrestrial Microbiology, Marburg 35043, Germany; Center for Synthetic Microbiology, Philipps-Universität Marburg, Marburg 35032, Germany

## Contributions

R.I and T.C contributed equally. R.I., T.C., T.J.E., and M.B. F.W, L.C.M.M. conceived, planned, and designed the study. R.I. T.C., M.B., C.K.B, L.A., J.B, P.S, J.B., M.K., performed the experiments and data analysis. F.R. performed the western-blot experiments. J.B. performed the fluorescence microscopy and data analysis. N.P. performed the metabolomics experiments and data analysis. T.G performed the proteomics experiments and data analysis. T.J.E., R.I. T.C., F.W. wrote and edited the manuscript with the input of all authors. T.J.E. acquired funding and supervised the work. All authors approved the final manuscript.

## Extended Data

**Extended Data Fig. 1:**
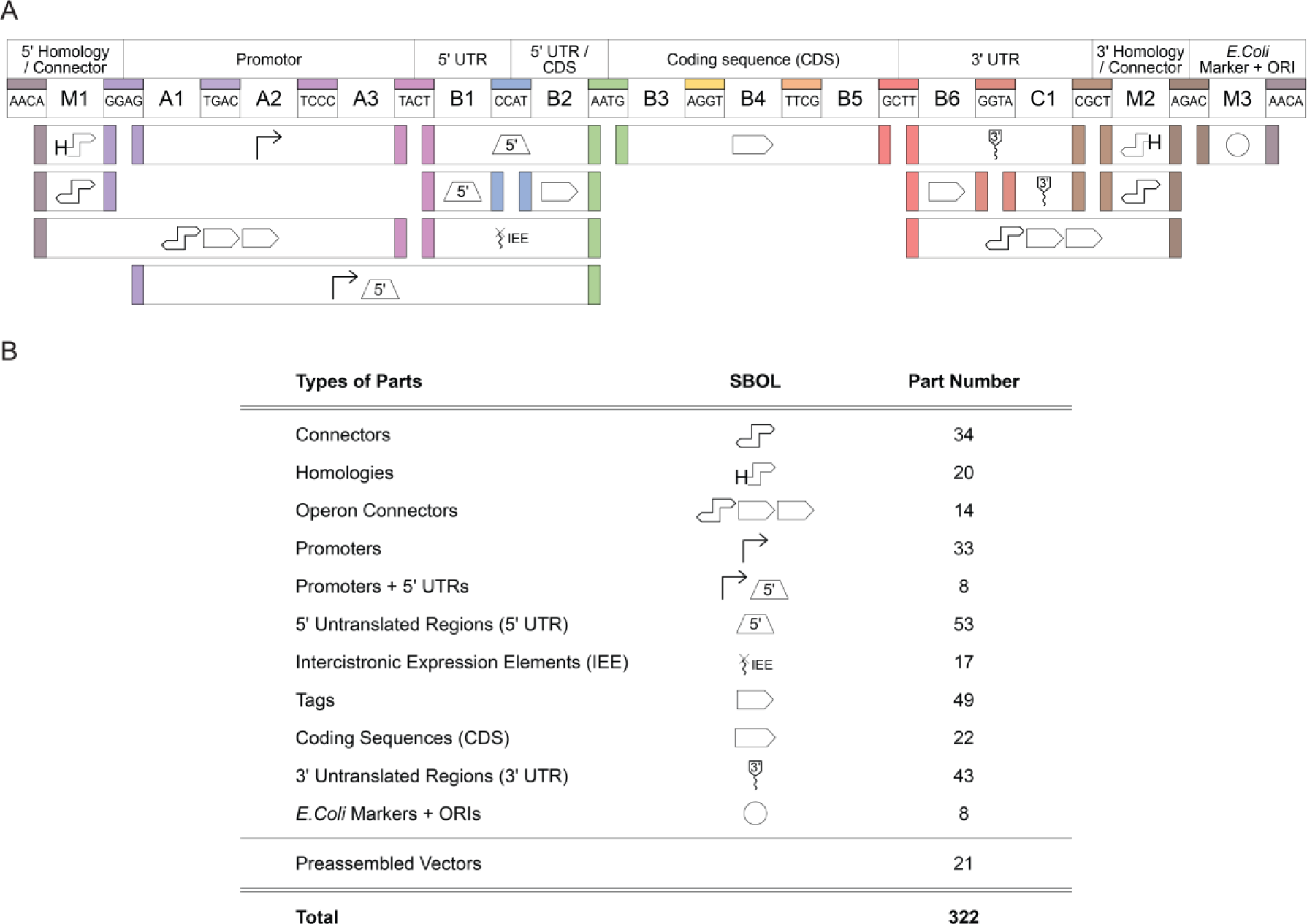
Architecture of the novel chloroplast modular cloning (MoClo) system and overview of the created genetic parts. **b,** The architecture follows the Phytobrick standard^47^ and conserves the cloning overhangs of this system, as well as the nomenclature for the part types (A1-C1). The position M1-3 allow for higher modularity as reported by Stukenberg *et al.* Number of parts are indicated for each position. **b**, List of parts. Each type of parts is listed, their SBOL depicted and the number of parts associated indicated.

**Extended Data Fig. 2:**
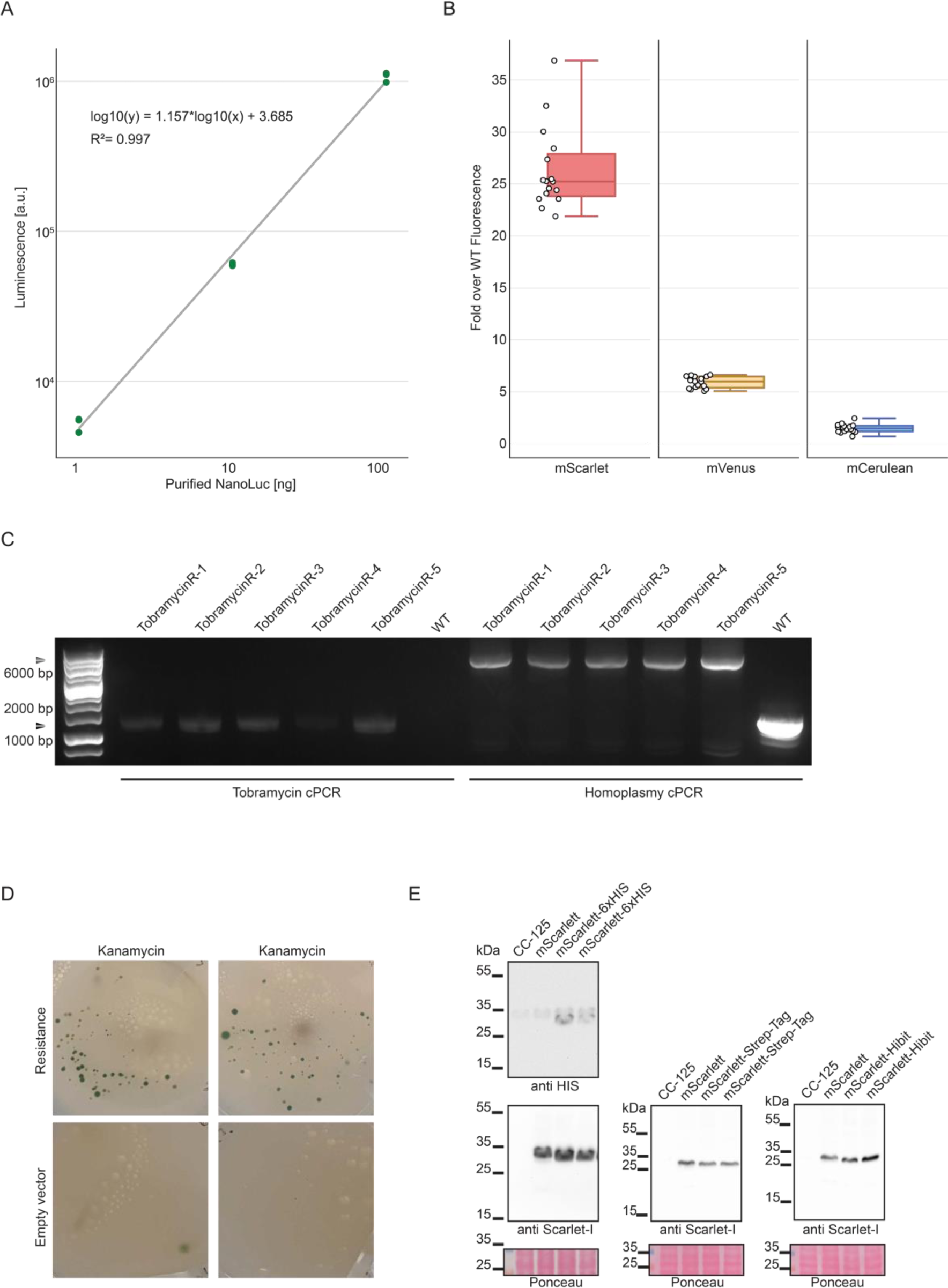
Reporters and selection markers. **a**, Calibration curve for purified NanoLuc concentration (ng) versus luminescence intensity (a.u.). The relationship between nanoluciferase abundance and luminescence is linear. **b**, Fluorescent reporter characterization. Fold over WT fluorescence is displayed for each strain, normalized by chlorophyll content, each expressing a different fluorescent reporter. n_biological_=16. **c,** PCR analysis of tobramycin-resistant strains. The first section of the gel illustrates PCR products of the tobramycin resistance gene, exclusive to the five genetically modified strains, with ∼1500 bp bands, absent in WT. The latter section shows PCR products from the insertion site, with ∼8000 bp bands in engineered strains and a 2000 bp band in WT, indicating homoplasmy in the modified strains. **d,** Selection marker for chloroplast transformation, with two replicates and two negative controls for each one. Successful selection after chloroplast transformation can be observed. **e,** Western blot analysis of transplastomic strains producing mScarlet HIS-tagged, Strep-tagged, Hibit-tagged in the Cterm position. Each strain is also tested for the presence of mScarlet using mScarlet specific probes. Ponceau gels are shown for each gel.

**Extended Data Fig. 3:**
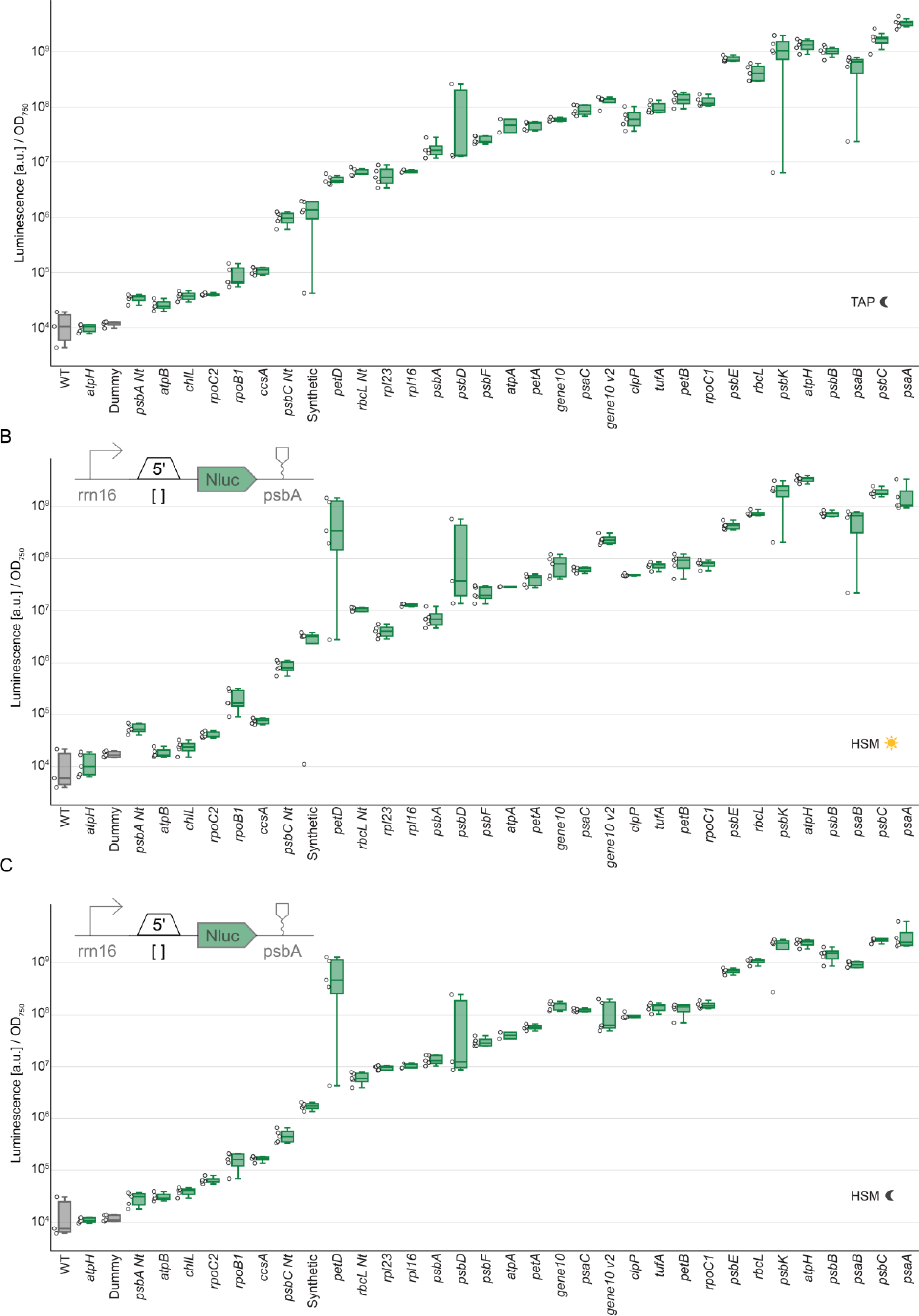
Characterization of 5’ UTR under different light and medium conditions. 5’UTRs were characterized by measuring NanoLuc activity of transplastomic strains containing different constructs, which differ in their 5’UTR. Nanoluc signal of a transplastomic strain containing a NanoLuc expression cassette (green) is plotted as arbitrary units [a.u.] normalized to OD and compared to WT strain (gray). For all measurements, n_biological_= 5, n_technical_= 3. **a,** NanoLuc assay under TAP dark conditions, **b,** under HSM light conditions and **c,** under HSM dark conditions. PetD stands out under HSM conditions.

**Extended Data Fig. 4:**
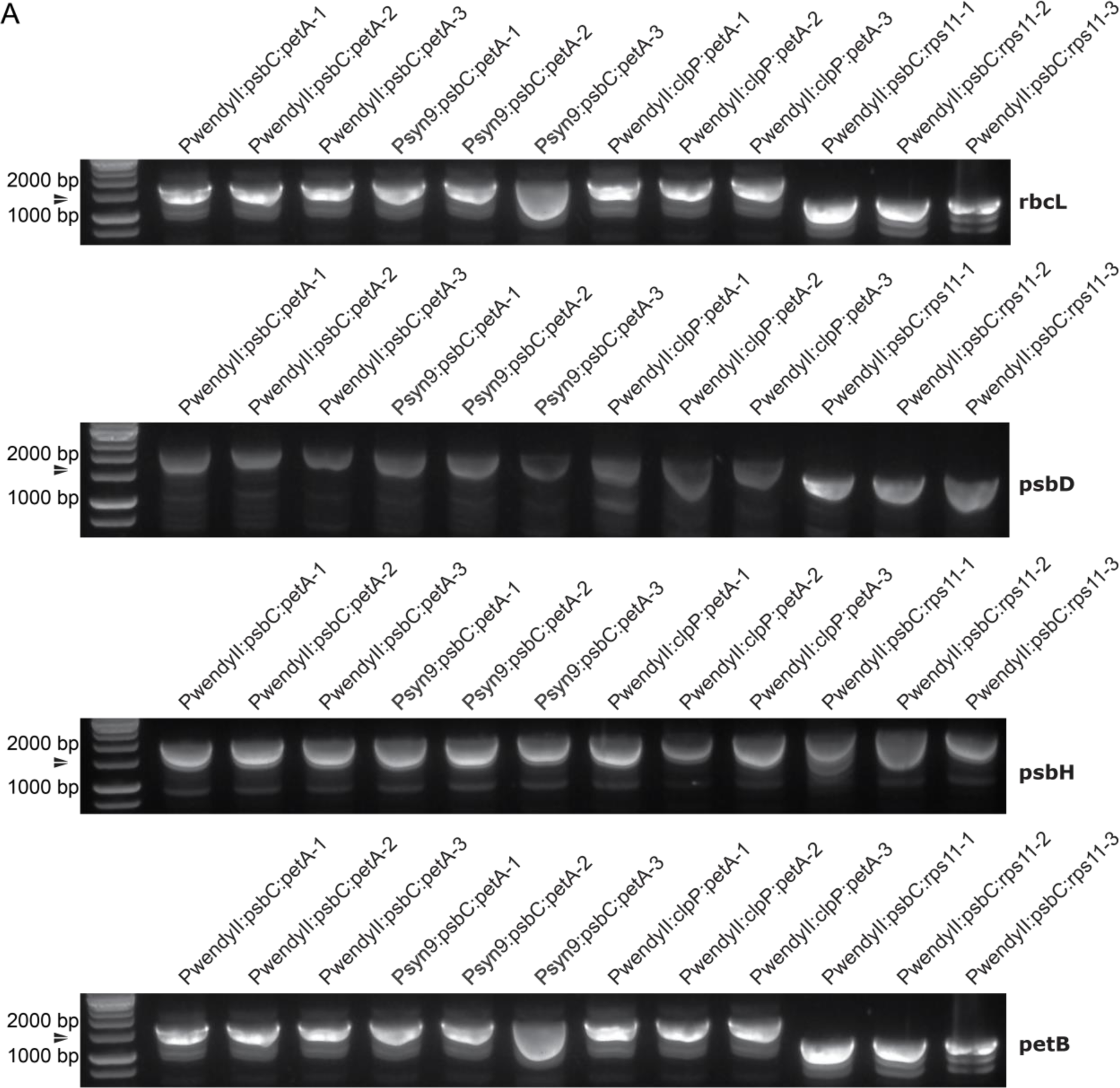
Genotyping of transplastomic strains. Gel depiction of cPCR amplification for the NanoLuc reporter gene across transplastomic strains shown in Figure 5c, with four distinct constructs integrated at separate sites in the genome. For each construct, n_biological_=3. A unique band, ranging from ∼1500-2000 bp, is visible in genetically modified strains across all four constructs and integration sites, but is not present in the WT.

**Extended Data Fig. 5:**
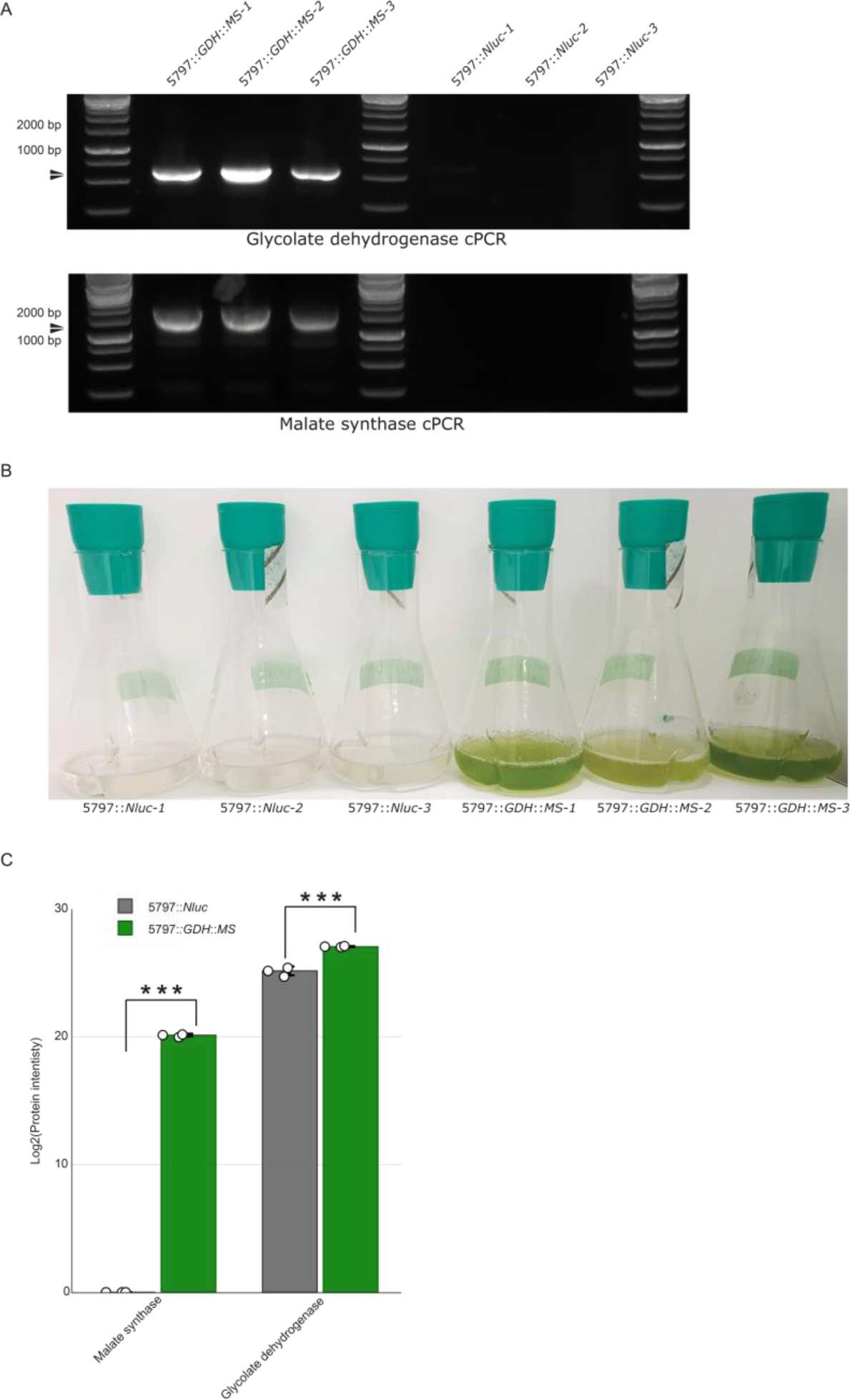
Genotyping and phenotyping of transplastomic strains containing the synthetic photorespiratory bypass. **a,** Gel depiction of cPCR amplification for the two heterologous genes glycolate dehydrogenase (GDH) and Malate synthase (MS) gene across transplastomic strains shown in Figure 6c-k. A unique band of ∼500 bp and 1500bp, for GDH and MS respectively, is visible in genetically modified strains but is not present in the control (5797::Nluc). For each experiment, n_biological_=3. **b,** Phenotypic comparison of strains expressing GDH and MS after 15 days. The image presents three biological replicates of the 5797::*GDH*::*MS* strain, demonstrating robust and dense growth. In contrast, control strains (5797::*Nluc*) exhibit no further growth. **c**. Proteomic analysis of the modified strains. The chart compares protein levels of MS and GDH in both the 5797::*GDH*::*MS* and 5797::*Nluc* strains, revealing significantly higher protein intensities in the 5797::*GDH*::*MS* strain for both proteins. Statistical significance is denoted by *** for a p-value < 0.001 (MS p-value < 0.000001, GDH p-value = 0.000623).

## Notes

### Competing Interest Statement

The authors have declared no competing interest.

### Summary of Updates

A spelling mistake in the name of one of the authors was corrected

