## Supplementary Information for "Advancing chloroplast synthetic biology through high-throughput plastome engineering of *Chlamydomonas reinhardtii*"

* Corresponding author

Inventory of Supplementary Information

**Extended Data Figures**	**3**

Extended Data Figure 1: Architecture of the novel chloroplast modular cloning (MoClo) system	3

Extended Data Figure 2: Reporters and selection markers	4

Extended Data Figure 3: Characterization of 5’ UTR under different light and medium conditions	5

Extended Data Figure 4: Genotyping of transplastomic strains	6

Extended Data Figure 5: Genotyping and phenotyping of transplastomic strains containing the synthetic photorespiratory bypass	7

**Supplementary Figures**	8

Supplementary Figure 1: Gel analysis of cPCR amplification for NanoLuc	8

Supplementary Figure 2: cPCR and sequencing for strain genomic characterization	9

**Supplementary Texts**	10

Supplementary text 1:Fluorescence microscopy settings for each fluorophore, chlorophyll and brightfield	10

Supplementary text 2: **:** PIXL colony detection parameters	12

Supplementary text 3: **:** FACS analysis settings	13

**Supplementary Tables**	14

Supplementary table 1: Mass Spectrometry parameters for CoAs	14

Supplementary table 2: Mass Spectrometry parameters for organic acids	14

Supplementary table 3: Mass Spectrometry parameters for amino acids	15

Supplementary table 4: Mass Spectrometry parameters for energy metabolites	16

### **Extended data Figures**

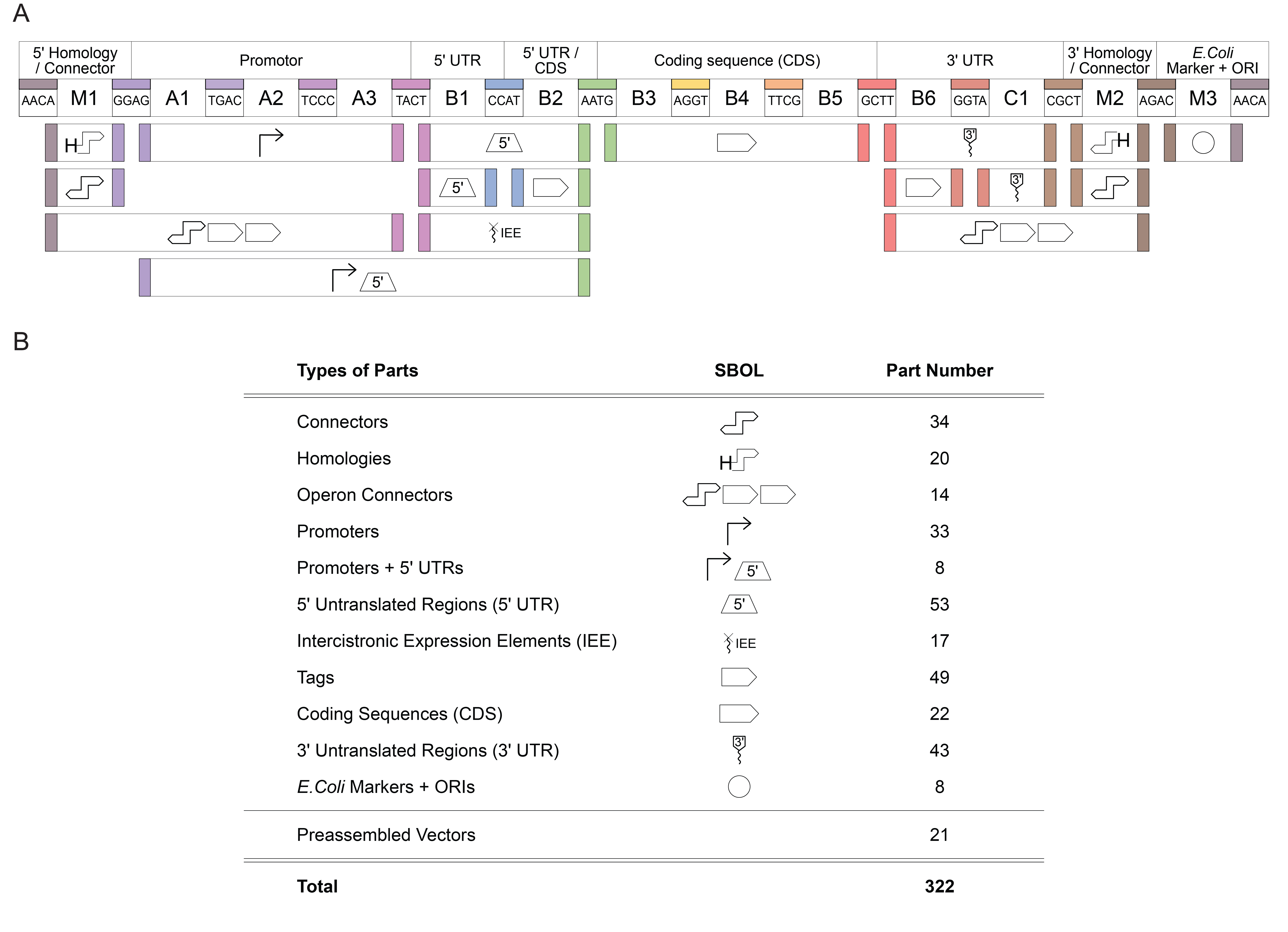

**Extended Data Fig.1:** Architecture of the novel chloroplast modular cloning (MoClo) system and overview of the created genetic parts. **b,** The architecture follows the Phytobrick standard^47^ and conserves the cloning overhangs of this system, as well as the nomenclature for the part types (A1-C1). The position M1-3 allow for higher modularity as reported by Stukenberg *et al.* Number of parts are indicates for each position. **b**, List of parts. Each type of parts is listed, their SBOL depicted and the number of parts associated indicated.

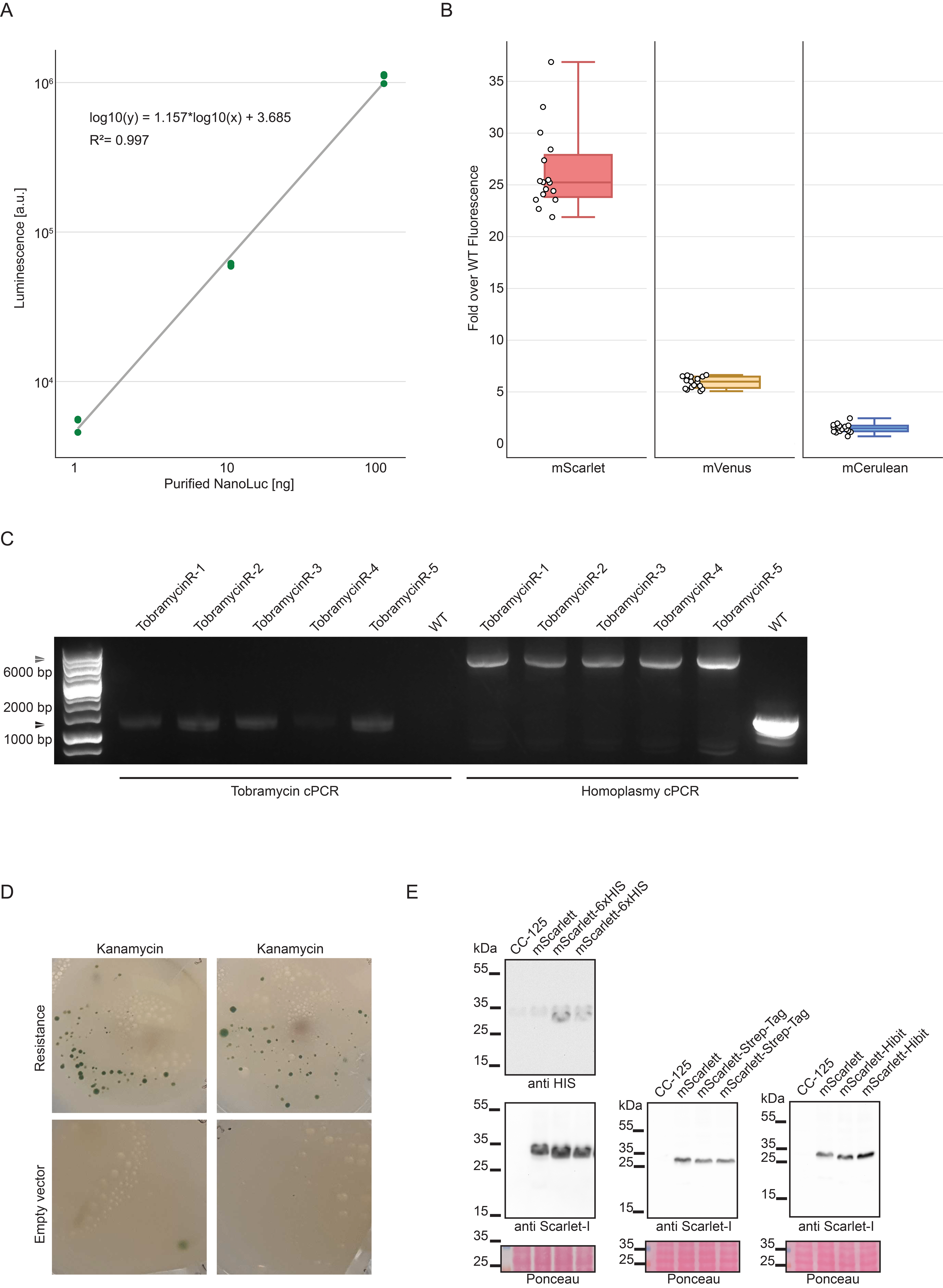

**Extended Data Fig.2: Reporters and selection markers**. **a**, Calibration curve for purified NanoLuc concentration (ng) versus luminescence intensity (a.u.). The relationship between nanoluciferase abundance and luminescence is linear. **b**, Fluorescent reporter characterization. Fold over WT fluorescence is displayed for each strain, normalized by chlorophyll content, each expressing a different fluorescent reporter. n_biological_=16. **c,** PCR analysis of tobramycin-resistant strains. The first section of the gel illustrates PCR products of the tobramycin resistance gene, exclusive to the five genetically modified strains, with ~1500 bp bands, absent in WT. The latter section shows PCR products from the insertion site, with ~8000 bp bands in engineered strains and a 2000 bp band in WT, indicating homoplasmy in the modified strains. **d,** Selection marker for chloroplast transformation, with two replicates and two negative controls for each one. Successful selection after chloroplast transformation can be observed. **e,** Western blot analysis of transplastomic strains producing mScarlet HIS-tagged, Strep-tagged, Hibit-tagged in the Cterm position. Each strain is also tested for the presence of mScarlet using mScarlet specific probes. Ponceau gels are shown for each gel.

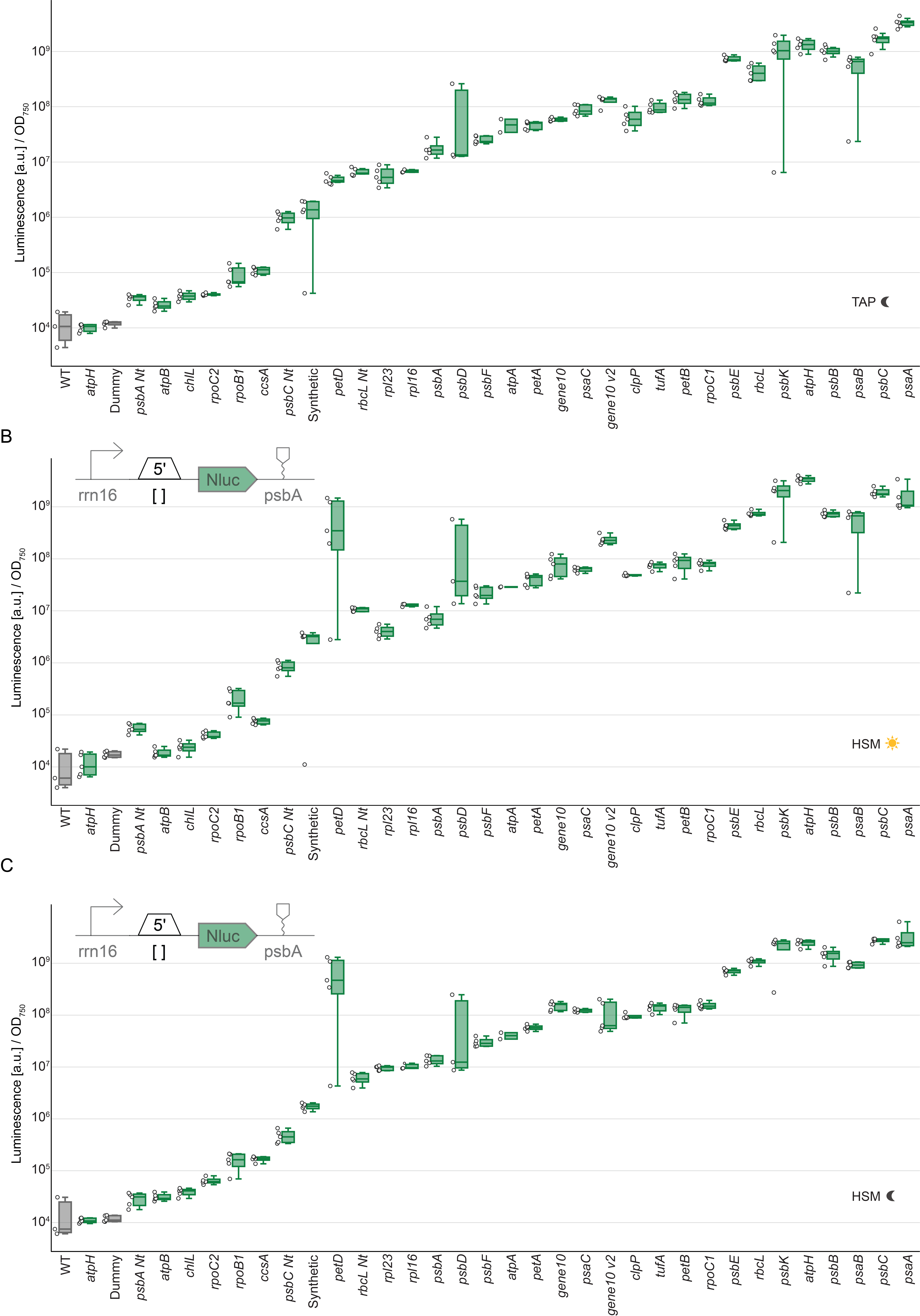

**Extended Data Fig3:** Characterization of 5’ UTR under different light and medium conditions.
5’UTRs were characterized by measuring NanoLuc activity of transplastomic strains containing different constructs, which differ in their 5’UTR. Nanoluc signal of a transplastomic strain containing a NanoLuc expression cassette (green) is plotted as arbitrary units [a.u.] normalized to OD and compared to WT strain (gray). For all measurements, n_biological_= 5, n_technical_= 3. **a,** NanoLuc assay under TAP dark conditions, **b,** under HSM light conditions and **c,** under HSM dark conditions. PetD stands out under HSM conditions.

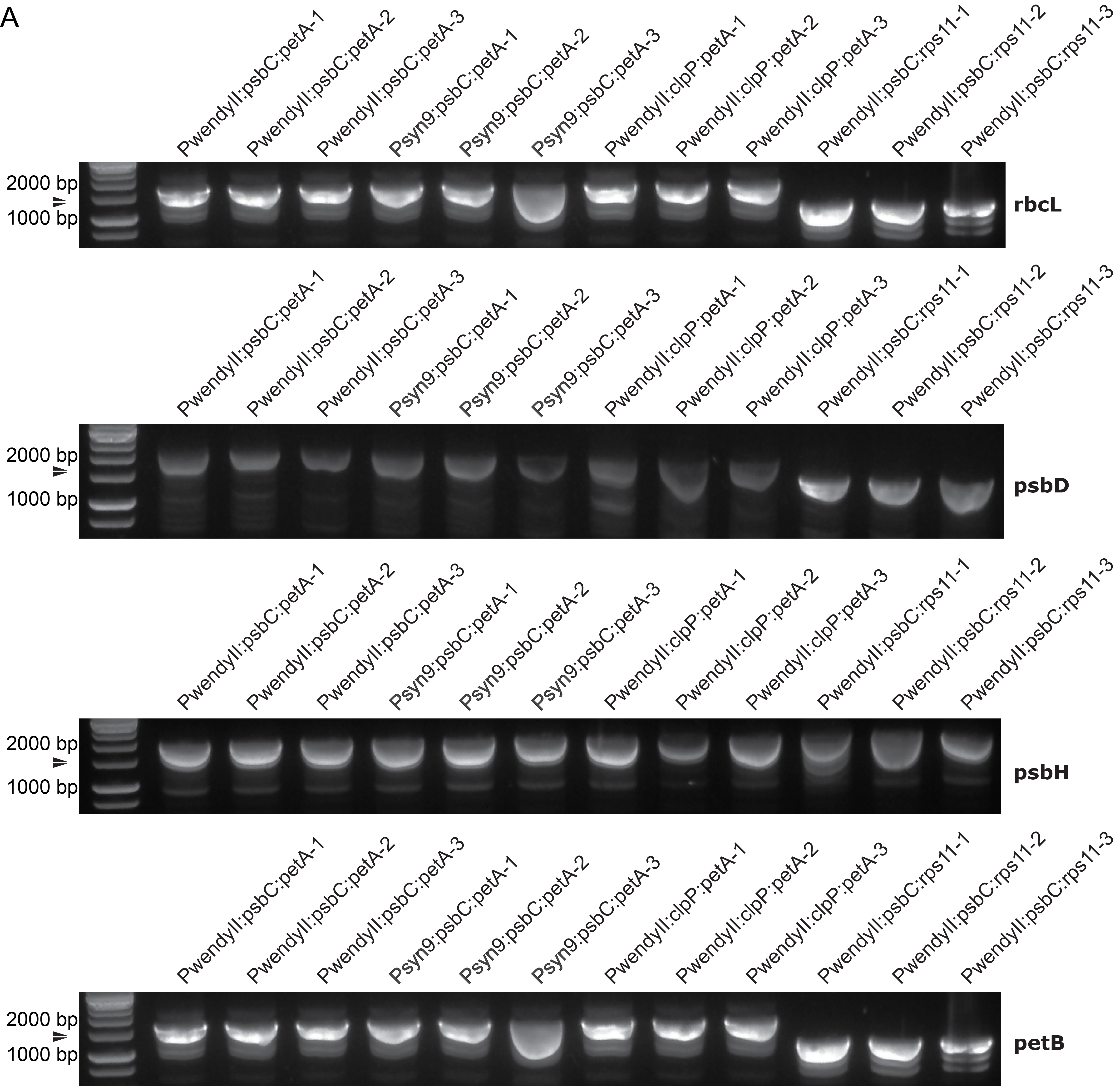

**Extended Data Fig.4:** Genotyping of transplastomic strains**.** Gel depiction of cPCR amplification for the NanoLuc reporter gene across transplastomic strains shown in Figure 5c, with four distinct constructs integrated at separate sites in the genome. For each construct, n_biological_=3. A unique band, ranging from ~1500-2000 bp, is visible in genetically modified strains across all four constructs and integration sites, but is not present in the WT.

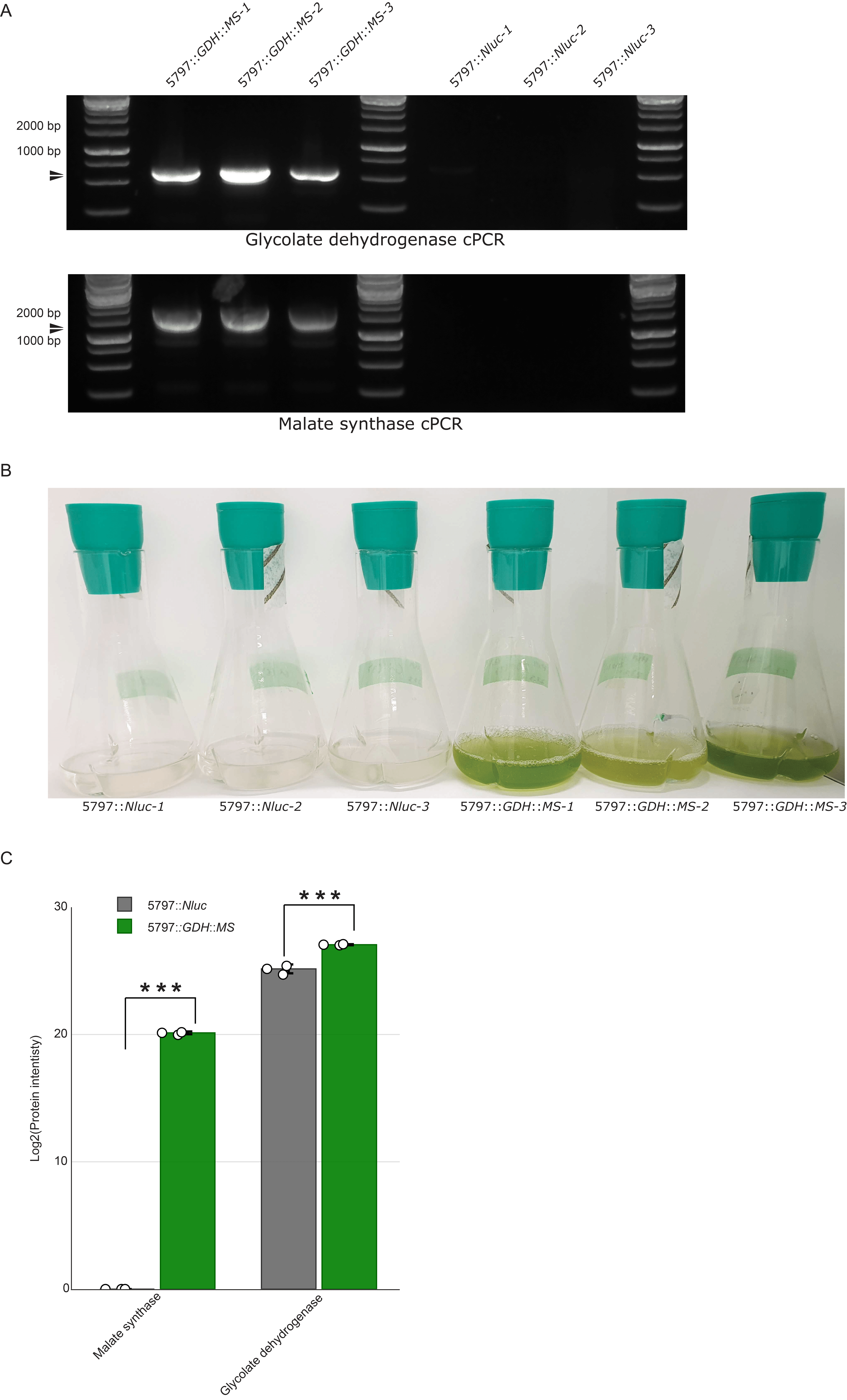

**Extended Data Fig.5:** Genotyping and phenotyping of transplastomic strains containing the synthetic photorespiratory bypass. **a,** Gel depiction of cPCR amplification for the two heterologous genes glycolate dehydrogenase (GDH) and Malate synthase (MS) gene across transplastomic strains shown in Figure 6c-k. A unique band of ~500 bp and 1500bp, for GDH and MS respectively, is visible in genetically modified strains but is not present in the control (5797::Nluc). For each experiment, n_biological_=3. **b,** Phenotypic comparison of strains expressing GDH and MS after 15 days. The image presents three biological replicates of the 5797::*GDH*::*MS* strain, demonstrating robust and dense growth. In contrast, control strains (5797::*Nluc*) exhibit no further growth. **c**. Proteomic analysis of the modified strains. The chart compares protein levels of MS and GDH in both the 5797::*GDH*::*MS* and 5797::*Nluc* strains, revealing significantly higher protein intensities in the 5797::*GDH*::*MS* strain for both proteins. Statistical significance is denoted by *** for a p-value < 0.001 (MS p-value < 0.000001, GDH p-value = 0.000623).

### **Supplementary Figures**

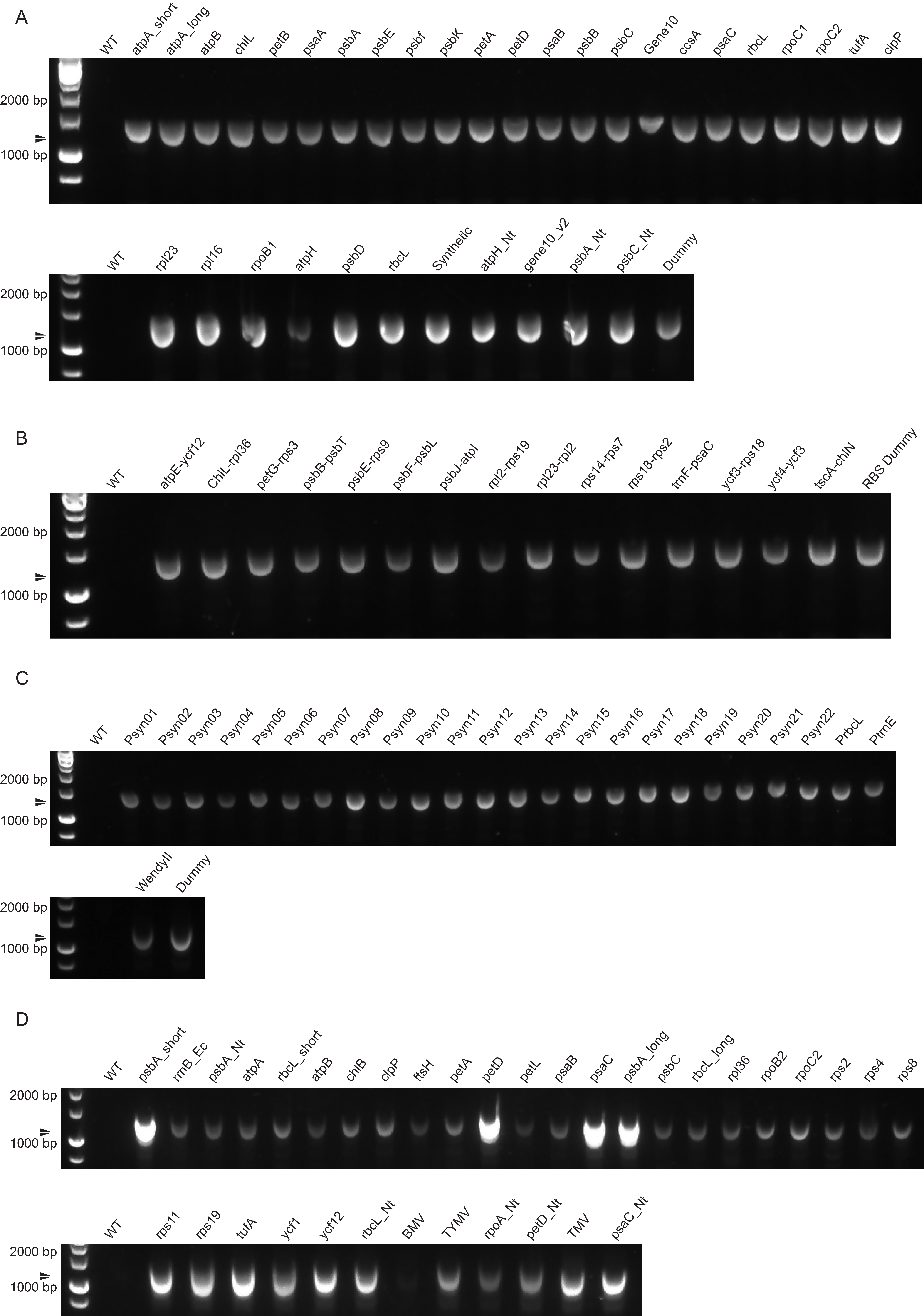

**Supplementary figure 1**: Gel analysis of cPCR amplification for the NanoLuc reporter gene in transplastomic strains, characterized in Figures 3-4, featuring one representative colony for each construct. A distinct band, approximately 1100-1300 bp in size, is observed in the genetically modified strains, absent in the WT. **a-d,** cPCR for NanoLuc in strains containing various 5’UTRs, IEEs, Promoters, and 3’UTRs respectively.

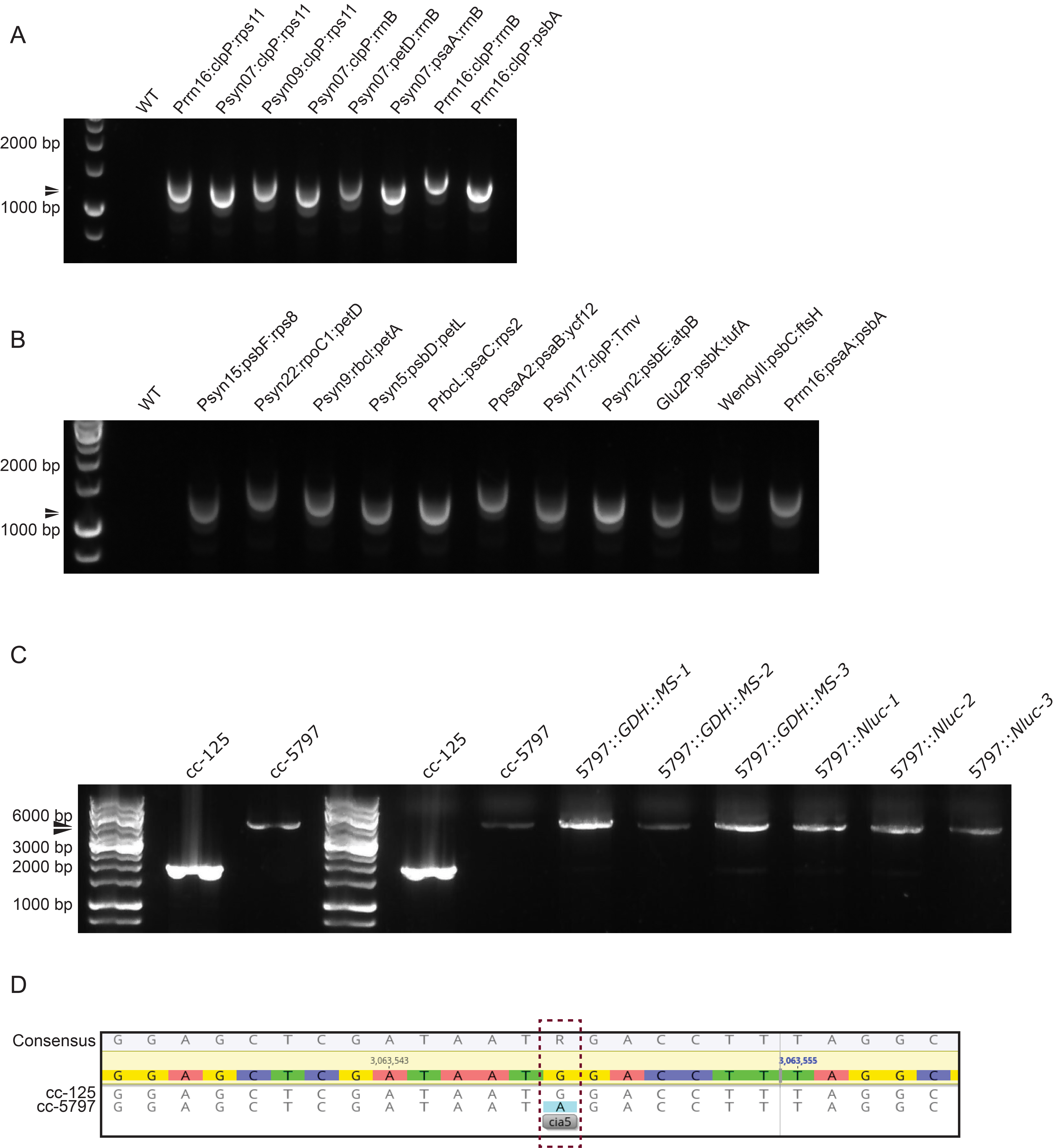

**Supplementary figure 2**: cPCR and sequencing for strain genomic characterization. **a-b,** Gel analysis of cPCR amplification for the NanoLuc reporter gene in transplastomic strains, characterized in Figures 5, featuring one representative colony for each construct. A distinct band, approximately 1100-1300 bp in size, is observed in the genetically modified strains, absent in the WT. **c** cPCR analysis of the glycolate dehydrogenase gene in cc-125 WT, cc-5797 mutant, and all engineered strains depicted in Figure 6. The transition from a 2000 bp band in cc-125 to a 6000 bp band in cc-5797, attributable to an insertion cassette. This band shift is consistently present in all engineered cc-5797 strains, verifying the insertion cassette's presence across these variants. **d,** Sequencing analysis reveals a point mutation at the cia5 locus in the nuclear genome of the cc-5797 mutant, absent in the cc-125 WT. A dark red dotted-lined box highlights this mutation, confirming the previously reported genotype.

### **Supplementary Text**

**Supplementary text 1:** Fluorescence microscopy settings for each fluorophore, chlorophyll and brightfield.

mCherry

Excitation laser:	561 nm

Laser power:	2.5%

Pinhole:	103

Imaging mode:	SUPERRES

Master gain:	750

Digital gain:	1.5

Main beam splitter (MBS):	488/561/633

Second beam splitter (SBS):	LP 570

Emission dual filters:	BP 420-480 + BP 495-620

mScarlet-I

Excitation laser:	561 nm

Laser power:	2.5%

Pinhole:	109

Imaging mode:	SUPERRES

Master gain:	750

Digital gain:	1.5

Main beam splitter (MBS):	488/561/633

Second beam splitter (SBS):	LP 570

Emission dual filters:	BP 420-480 + BP 495-620

mVenus

Excitation laser:	514 nm

Laser power:	5%

Pinhole:	103

Imaging mode:	SUPERRES

Master gain:	775

Digital gain:	1.0

Main beam splitter (MBS):	458/514

Second beam splitter (SBS):	LP 525

Emission dual filters:	BP 420-480 + BP 495-550

mCerulean

Excitation laser:	458 nm

Laser power:	5%

Pinhole:	103

Imaging mode:	SUPERRES

Master gain:	775

Digital gain:	1.0

Main beam splitter (MBS):	458/561

Second beam splitter (SBS):	SP 615

Emission dual filters:	BP 465-505 + LP 525

Chlorophyll

Excitation laser:	633 nm

Laser power:	1%

Pinhole:	103

Imaging mode:	SUPERRES

Master gain:	720

Digital gain:	1.0

Main beam splitter (MBS):	488/561/633

Second beam splitter (SBS):	BP 570-625

Emission dual filters:	BP 570-620 + LP 645

Brightfield

Excitation laser:	561 nm

Laser power:	2%

Pinhole:	76.4

Imaging mode:	T-PMT

Master gain:	150

Digital gain:	1.0

**Supplementary text 2:** PIXL colony detection parameters. Imaging settings and algorithm selection.

Mode:	White

Gain:	39.4%

Gamma:	39.4%

Exposure:	10ms

Saturation:	34.4

White Balance - Red:	34.4%

White Balance - Green:	25%

White Balance - Blue:	74.3

Focus:	Auto

Lighting power:	70%

Algorithm:	Colony Separation

Organism:	S. cerevisiae (dark colonies)

Blueness Filter:	0 - 1

Circularity Filter:	0 - 1

Greenness Filter:	0 - 1

Intensity Filter:	0 - 1

Proximity Filter:	0 - 100

Radius Filter:	0.25 – 2.5

Redness Filter:	0 - 1

**Supplementary text 3:** FACS analysis settings.

Laser (mScarlet-I):	561 nm

Filter (mScarlet-I):	570-630 nm

Filter Gain (mScarlet-I):	40%

Laser (Chlorophyll):	638 nm

Filter (Chlorophyll):	690-750 nm

Filter Gain (Chlorophyll):	40%

### **Supplementary Tables**

**Supplementary Table 1**: Mass Spectrometry parameters for CoAs.

| Name | Precurser Ion | Product Ion | Collision energy [V] | Fragmentor Voltage [V] | Cell Accelorator Voltage [V] | Dwell time [msec] | Polarity |
| --- | --- | --- | --- | --- | --- | --- | --- |
| CoA | 768.12 | 428  261.1 | 35  35 | 380  380 | 5  5 | 220  220 | Positive  Positive |
| ACoA | 810.1 | 428  302.2 | 31  31 | 380  380 | 5  5 | 220  220 | Positive  Positive |

**Supplementary Table 2**: Mass Spectrometry parameters for organic acids.

| Name | Precurser Ion | Product Ion | Collision energy [V] | Fragmentor Voltage [V] | Cell Accelorator Voltage [V] | Dwell time [msec] | Polarity |
| --- | --- | --- | --- | --- | --- | --- | --- |
| Citrate | 191 | 111.1  85.1 | 11  14 | 380  380 | 5  5 | 20  20 | Negative |
| Alphaketoglutarate | 145.1 | 101.1  57.2 | 5  8 | 380  380 | 5  5 | 20  20 | Negative |
| Malate | 133.1 | 115.1  71.2 | 8  14 | 380  380 | 5  5 | 20  20 | Negative |
| Succinate | 117.2 | 73.2  55.1 | 9  15 | 380  380 | 5  5 | 20  20 | Negative |
| Fumarate | 115.1 | 71.2  27.3 | 4  9 | 380  380 | 5  5 | 20  20 | Negative |
| Lactate | 89.2 | 89.2  71.3 | 0  10 | 380  380 | 5  5 | 20  20 | Negative |
| Pyruvate | 87.1 | 87.1  43.1 | 0  4 | 380  380 | 5  5 | 20  20 | Negative |
| Glycolate | 75.2 | 75.2  47.2 | 0  6 | 380  380 | 5  5 | 20  20 | Negative |
| Glyoxylate | 73.2 | 73.2  45.2 | 0  7 | 380  380 | 5  5 | 20  20 | Negative |

**Supplementary Table 3**: Mass Spectrometry parameters for amino acids.

| Compound | Precurser | Product | Dwell time  [msec] | Fragmenter Voltage  [V] | Collision Energy  [V] | Cell Accelerator Voltage [V] | Polarity |
| --- | --- | --- | --- | --- | --- | --- | --- |
| Tryptophane | 205.1 | 188  145.9 | 20  20 | 380  380 | 7  17 | 5  5 | Positive  Positive |
| Tyrosine | 182.1 | 165  136.1 | 20  20 | 380  380 | 6  12 | 5  5 | Positive  Positive |
| Arginie | 174.9 | 116  70.2 | 20  20 | 380  380 | 12  29 | 5  5 | Positive  Positive |
| Phenylalanine | 166.1 | 120.2  103.1 | 20  20 | 380  380 | 13  32 | 5  5 | Positive  Positive |
| Histidine | 156.1 | 110.1  83 | 20  20 | 380  380 | 16  30 | 5  5 | Positive  Positive |
| Methionine | 150.1 | 133  104 | 20  20 | 380  380 | 7  7 | 5  5 | Positive  Positive |
| Glutamate | 148.1 | 84.1  56.1 | 20  20 | 380  380 | 17  34 | 5  5 | Positive  Positive |
| Glutamine | 147.2 | 130.1  84.2 | 20  20 | 380  380 | 8  17 | 5  5 | Positive  Positive |
| Lysine | 147.1 | 130.1  84.1 | 20  20 | 380  380 | 8  19 | 5  5 | Positive  Positive |
| Aspertate | 134.1 | 88  74 | 20  20 | 380  380 | 9  14 | 5  5 | Positive  Positive |
| Asparagine | 133. | 87.1  74.2 | 20  20 | 380  380 | 17  16 | 5  5 | Positive  Positive |
| Isoleucine | 132.1 | 86.1  69.1 | 20  20 | 380  380 | 8  18 | 5  5 | Positive  Positive |
| Leucine | 132.1 | 86.1  30.3 | 20  20 | 380  380 | 8  18 | 5  5 | Positive  Positive |
| Threonine | 120.2 | 74.1  55.9 | 20  20 | 380  380 | 8  18 | 5  5 | Positive  Positive |
| Valine | 118.1 | 72  55.1 | 20  20 | 380  380 | 9  23 | 5  5 | Positive  Positive |
| Proline | 116 | 70  43.3 | 20  20 | 380  380 | 15  35 | 5  5 | Positive  Positive |
| Serine | 106.1 | 60.2  42.2 | 20  20 | 380  380 | 12  11 | 5  5 | Positive  Positive |
| Alanine | 90 | 44.1 | 20 | 380 | 12 | 5 | Positive |
| Glycine | 76.1 | 30.3  28.3 | 20  20 | 380  380 | 32  32 | 5  5 | Positive  Positive |

**Supplementary Table 4**: Mass Spectrometry parameters for energy metabolites.

| Name | Precurser Ion | Product Ion | Collision energy [V] | Fragmentor Voltage [V] | Cell Accelorator Voltage [V] | Dwell time [msec] | Polarity |
| --- | --- | --- | --- | --- | --- | --- | --- |
| NAD | 664.1 | 524  428 | 18  26 | 380  380 | 5  5 | 50  50 | Positive  Positive |
| NADH | 666.1 | 649  514 | 17  23 | 380  380 | 5  5 | 50  50 | Positive  Positive |
| NADP | 744.1 | 604  508 | 20  32 | 380  380 | 5  5 | 50  50 | Positive  Positive |
| NADPH | 746.1 | 729  135.8 | 17  40 | 380  380 | 5  5 | 50  50 | Positive  Positive |
| AMP | 346 | 96  78.9 | 24  30 | 380  380 | 5  5 | 50  50 | Negative  Negative |
| ADP | 425.9 | 327.8  133.9 | 17  22 | 380  380 | 5  5 | 50  50 | Negative  Negative |
| ATP | 505.9 | 407.9  158.8 | 21  28 | 380  380 | 5  5 | 50  50 | Negative  Negative |
